# ZCWPW1 is recruited to recombination hotspots by PRDM9, and is essential for meiotic double strand break repair

**DOI:** 10.1101/821678

**Authors:** Daniel Wells, Emmanuelle Bitoun, Daniela Moralli, Gang Zhang, Anjali Gupta Hinch, Peter Donnelly, Catherine Green, Simon R Myers

## Abstract

During meiosis, homologous chromosomes pair (synapse) and recombine, enabling balanced segregation and generating genetic diversity. In many vertebrates, recombination initiates with double-strand breaks (DSBs) within hotspots where PRDM9 binds, and deposits H3K4me3 and H3K36me3. However, no protein(s) recognising this unique combination of histone marks have yet been identified.

We identified *Zcwpw1*, which possesses H3K4me3 and H3K36me3 recognition domains, as highly co-expressed with *Prdm9*. Here, we show that ZCWPW1 has co-evolved with PRDM9 and, in human cells, is strongly and specifically recruited to PRDM9 binding sites, with higher affinity than sites possessing H3K4me3 alone. Surprisingly, ZCWPW1 also recognizes CpG dinucleotides, including within many Alu transposons.

Male *Zcwpw1* homozygous knockout mice show completely normal DSB positioning, but persistent DMC1 foci at many hotspots, particularly those more strongly bound by PRDM9, severe DSB repair and synapsis defects, and downstream sterility. Our findings suggest a model where ZCWPW1 recognition of PRDM9-bound sites on either the homologous, or broken, chromosome is critical for synapsis, and hence fertility.

**Graphical Abstract Legend:** In humans and other species, recombination is initiated by double strand breaks at sites bound by PRDM9. Upon binding, PRDM9 deposits the histone marks H3K4me3 and H3K36me, but the functional importance of these marks has remained unknown. Here, we show that PRDM9 recruits ZCWPW1, a reader of both these marks, to its binding sites genome-wide. ZCWPW1 does not help position the breaks themselves, but is essential for their downstream repair and chromosome pairing, and ultimately meiotic success and fertility in mice.

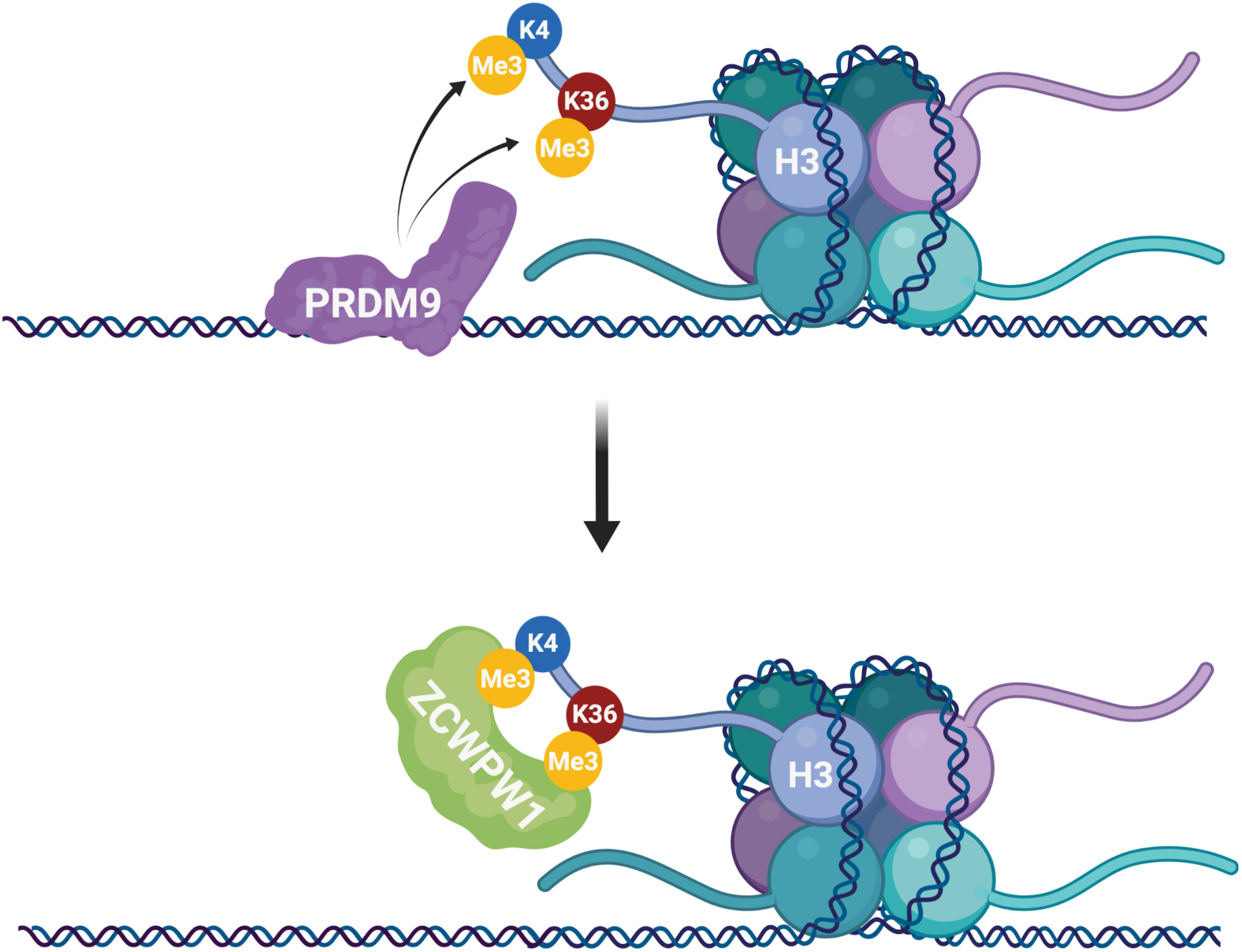

## Introduction

Meiosis is a specialised cell division, producing haploid gametes essential for reproduction. Uniquely during this process, homologous maternal and paternal chromosomes pair and exchange DNA (recombine) before undergoing balanced independent segregation. Alongside *de novo* mutations, this generates all genetic diversity - providing a substrate on which natural selection can act.

In humans, mice, and likely many other vertebrates (Baker et al., 2017), the locations at which recombination occurs is determined by the binding locations of PRDM9, which recognises a specific sequence motif encoded by its zinc finger array (Baudat et al., 2010; Myers et al., 2010; Parvanov et al., 2010). At these sites, Spo11 catalyzes recombination-initiating double strand breaks (DSBs), a subset of which are repaired by recombination with the homologous chromosome. The sister chromatid can also provide a repair substrate, e.g. for DSBs on the X chromosome in males, and there is evidence that this also occurs at some autosomal hotspots (Allen and Latt, 1976; R. Li et al., 2019; Lu and Yu, 2015).

In *Prdm9*-null mice, DSBs form at promoter regions rich in H3K4me3 (Brick et al., 2012), and fail to repair resulting in severe asynapsis, meiotic arrest at pachytene, and sterility in both males and females on the B6 background (Hayashi et al., 2005). We previously showed (Davies et al., 2016) that across DSB sites, increased binding of PRDM9 to the homologous chromosome aids synapsis and fertility in hybrid mice, implying an impact of binding downstream of DSB positioning. Mice where most DSBs occur at sites where PRDM9 does *not* bind the homologue show widespread asynapsis (Davies et al., 2016; Gregorova et al., 2018). More recently, we showed that binding of PRDM9 to the homologous chromosome promotes repair of DSBs by homologue-templated recombination, potentially by aiding homology search (R. Li et al., 2019).

DSB sites are processed by resection, resulting in single-stranded DNA (ssDNA) that becomes decorated with DMC1 (Hong et al., 2001; Neale and Keeney, 2006; Sehorn et al., 2004). In WT mice, DMC1 foci start to appear in early zygotene cells upon loading to the ssDNA ends of the DSBs created during leptotene. From mid-zygotene to early pachytene, as part of the recombinational repair process, DMC1 dissociates from the ssDNA and counts decrease until all breaks (except those on the XY chromosomes) are repaired in late pachytene (Moens et al., 2002). At DSB sites where the homologous chromosome is not bound by PRDM9, DMC1 signal is strongly elevated (Davies et al., 2016), suggesting delayed DSB repair, while fewer homologous recombination events occur (Hinch et al., 2019; R. Li et al., 2019), suggesting that (eventual) DSB repair may sometimes use a sister chromosome pathway. However, the underlying mechanism(s) by which PRDM9 effectively contributes to DSB repair are not yet known.

PRDM9 deposits both H3K4me3 and H3K36me3 histone methylation marks at the sites it binds, and this methyltransferase activity is essential for its role in DSB positioning (Diagouraga et al., 2018; Powers et al., 2016). What reads this unique combination of marks at recombination sites, however, is currently unknown. Notably, outside of hotspots and the pseudoautosomal region (PAR) on sex chromosomes, H3K4me3 and H3K36me3 occur at largely nonoverlapping locations (Powers et al., 2016), suggesting potentially highly specialised reader(s). Indeed, in somatic cells H3K4me3 is deposited mainly at promoters, in particular by the SET1 complex targeted by CXXC1/CFP1 and Wdr82 which binds Ser5 phosphorylated polymerase II (Barski et al., 2007; Lee and Skalnik, 2008, 2005). H3K36me3 is deposited by different methlytransferases, including SETD2 bound to Ser2 phosphorylated (elongating) polymerase II, and is enriched at exon bound nucleosomes, particularly for 3’ exons (reviewed in (McDaniel and Strahl, 2017; Wagner and Carpenter, 2012)). H3K36me3 has multiple important roles, including in directing DNA methylation by recruiting DNMT3B, somatic DSB repair by homologous recombination (Aymard et al., 2014; Carvalho et al., 2014; Pfister et al., 2014), mismatch repair by recruiting MSH6 (Huang et al., 2018; Li et al., 2013), and V(D)J recombination during lymphopoiesis (Ji et al., 2019).

Using single cell RNA-sequencing of mouse testis, we identified a set of genes co-expressed in (pre)leptotene cells which are highly enriched for genes involved in meiotic recombination (Jung et al., 2019). *Zcwpw1*, which ranks 3^rd^ in this set after *Prdm9* (2^nd^), is of unknown function but contains two recognised protein domains: CW and PWWP, shown to individually bind H3K4me3 and H3K36me3 respectively (F. He et al., 2010; Rona et al., 2016). This raises the attractive possibility that ZCWPW1 might recognize and physically associate with the same marks deposited by PRDM9 (Jung et al., 2019).

In humans, *ZCWPW1* is specifically expressed in testis (Carithers et al., 2015; Uhlén et al., 2015) (**Supplementary Figure 1**). It is also one of 104 genes specific to meiotic prophase in murine fetal ovary (Soh et al., 2015), further suggesting a conserved meiotic function. This was confirmed by a recent study showing that ZCWPW1 is required for male fertility, with ZCWPW1 hypothesised to recruit the DSB machinery to hotspot sites (M. Li et al., 2019). Here, we show that ZCWPW1 co-evolves with PRDM9, and is recruited to recombination hotspots by the combination of histone marks deposited by PRDM9. However, ZCWPW1 is *not* required for the positioning of DSBs at PRDM9-bound sites, which occurs normally in ZCWPW1-null mice. Instead, ZCWPW1 is required for proper inter-homologue interactions: synapsis and the repair of DSBs. In ZCWPW1-null mice, DMC1 signals show strong perturbations, with signals at autosomal hotspots resembling those on the X-chromosome, which does not have a homologue. Thus, ZCWPW1 represents the first protein directly positioned by PRDM9 binding, but impacting homologous DSB repair.

## Results

### *ZCWPW1* co-evolves with *PRDM9*

Based on publicly available databases, we identified likely ZCWPW1 orthologues in 167 species in total, aligned each to the human reference ZCWPW1 protein, and compared against a previous analysis of PRDM9 (Baker et al., 2017). The alignment reveals the regions containing the CW and PWWP domains to be the most conserved among species (**Figure 1**), while other parts of the protein appear to have been lost in some species. In addition, there is a region of moderate conservation downstream of the PWWP domain, not overlapping any known domain. Notably, an SYCP1 (SCP1) domain is annotated in the mouse protein only, which although only suggestive, is interesting given that SYCP1 physically connects homologous chromosomes in meiosis. In addition, protein threading suggests that the C terminal end of ZCWPW1 may contain a methyl-CpG binding domain (Methods, (Lobley et al., 2009).

**Figure 1.**
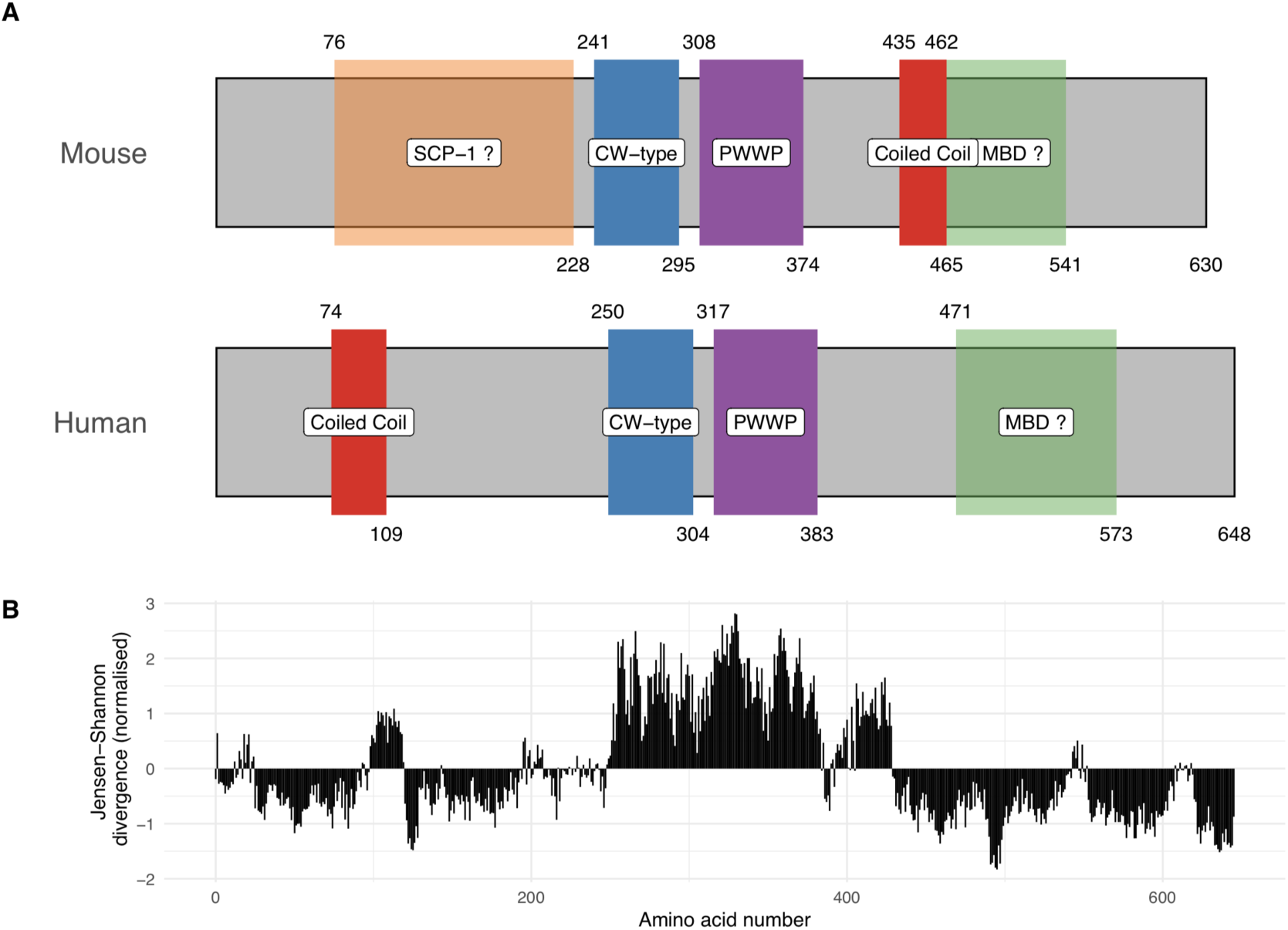
Domain organisation (A) and evolutionary conservation (B) of ZCWPW1. **(A)** Protein domains in the human and mouse proteins (source: UniProt). Start and end positions of each domain are shown above and below the rectangles respectively. Prediction of SCP-1 domain from (Marchler-Bauer and Bryant, 2004) and of MBDs from (Lobley et al., 2009) (Methods) (**B)** Conservation of human amino acids, normalised Jensen-Shanon divergence from (Capra and Singh, 2007; Johansson and Toh, 2010) using multiple alignment of 167 orthologues (Methods).

Because of incomplete available DNA and protein sequences, we are almost certain to miss some species where ZCWPW1 is present, and similarly it is challenging to identify PRDM9 orthologues (Baker et al., 2017). Despite this, we see extremely high overlap between PRDM9 and ZCWPW1 occurrence. All identified ZCWPW1 orthologues are in vertebrates, similar to identified PRDM9 orthologues (Baker et al., 2017). 99 of 167 (149) species with ZCWPW1 (PRDM9) respectively have the other protein; an even higher fraction of species have a close relative with an identified orthologue: 131 species with ZCWPW1 are in a family possessing PRDM9. Only 8 species possess ZCWPW1 but are in an *order* with no identified species possessing PRDM9: three amphibians including Xenopus frogs, and five placental mammals. Given widespread conservation of PRDM9 among mammals other than canids (Baker et al., 2017), these five might plausibly represent false negatives. Thus, ZCWPW1 appears to mainly occur in species, and even more clearly in groups of species, possessing PRDM9.

We identified ZCWPW1 orthologues across the full spectrum of vertebrates possessing PRDM9, including jawless, cartilaginous and bony fish, coelacanths, turtles, snakes and lizards, and mammals. Previous work (Baker et al., 2017) has identified at least 6 independent complete PRDM9 loss events; in birds and crocodiles; in three distinct groups of teleost fish (although these fish possess a PRDM9 “beta” orthologue with mutations in key catalytic amino acids within the SET domain); in canids; and tentatively, in amphibians. The first four losses appear to have been completely mirrored by corresponding ZCWPW1 losses, with no species previously studied by Baker *et al.* in these groups possessing a ZCWPW1 orthologue. For the latter two, we did find potential orthologues (e.g. in canids), but with mutations at positions in ZCWPW1 that are conserved among all PRDM9-SET-possessing species (Methods). Thus, there has been similar (co)evolution of presence/absence of ZCWPW1 and PRDM9.

We observe a greater number of species - 28 - possessing PRDM9 orthologues but having no relative closer than their class possessing an identified ZCWPW1 orthologue. All of these are bony fish. Strikingly, 24 of these 28 orthologues possess one or more mutations in their SET domain (**Supplementary Table 1**), predicted to disrupt H3K4me3 and/or H3K36me3 deposition. In strong contrast, 94% of PRDM9-containing species with non-mutant SET domains have a closer relative possessing ZCWPW1 (odds ratio=90; indicative p<10^−15^ by FET, though we note these observations are not all independent). This implies that ZCWPW1 is most often lost, whenever mutations occur in the SET domain of PRDM9. Interestingly, PRDM9-containing species with non-mutant SET domains are near-identical to those also possessing SSXRD domains, and SSXRD has been reported as essential for PRDM9’s H3K4me3 methyltransferase activity at hotspots *ex vivo* (Thibault-Sennett et al., 2018). However, other domains of PRDM9 (KRAB, and the zinc finger array) have been lost across multiple species, and it is hypothesized PRDM9 does not position recombination in these species (Baker et al., 2017). Nonetheless, they mainly retain ZCWPW1 (**Supplementary Table 1**). In conclusion, among vertebrates at least, it appears the set of species with ZCWPW1 is extremely similar to those species possessing PRDM9 with both an intact SET domain, and an SSXRD domain, with considerable evidence of co-evolution of gain/loss events for each protein. We note that this pattern is precisely what would be expected *a priori*, if ZCWPW1’s main function involves recognition of the histone modifications catalysed by PRDM9-SET during early meiosis, as predicted from functional considerations.

### Localisation of ZCWPW1 in meiosis and details of asynapsis in infertile male Zcwpw1^−/−^ mice

To investigate the role of ZCWPW1 during meiosis *in vivo*, we produced an antibody against the full-length recombinant mouse protein (**Supplementary Figure 2**) and studied the phenotype of a newly generated knockout (KO) mouse line for *Zcwpw1*, with a particular focus on fertility and meiotic recombination.

In testes from wild-type (WT) mice, we observe a dynamic localisation of ZCWPW1 protein (**Figure 2**), similar but non-identical to that reported in a recent study (M. Li et al., 2019). ZCWPW1 shows a strong, punctate nuclear staining excluding the pericentromeric regions (clustered into chromocenters brightly stained with DAPI) in zygotene and early pachytene cells; we detected transcript expression in earlier pre-leptotene to leptotene cells (Jung et al., 2019). In pachytene cells, ZCWPW1 expression drops, with the protein now mainly localised in the XY body and as bright foci at the ends of the synaptonemal complex labelled by SYCP3, not previously observed using an antibody raised against a 174 base-pair (bp) C-terminal region of the protein (M. Li et al., 2019). By diplotene, little expression is visible. Using FISH to label telomeric ends of chromosomes, we established that these discrete foci of ZCWPW1 are located at subtelomeric regions (**Supplementary Figure 3**).

**Figure 2.**
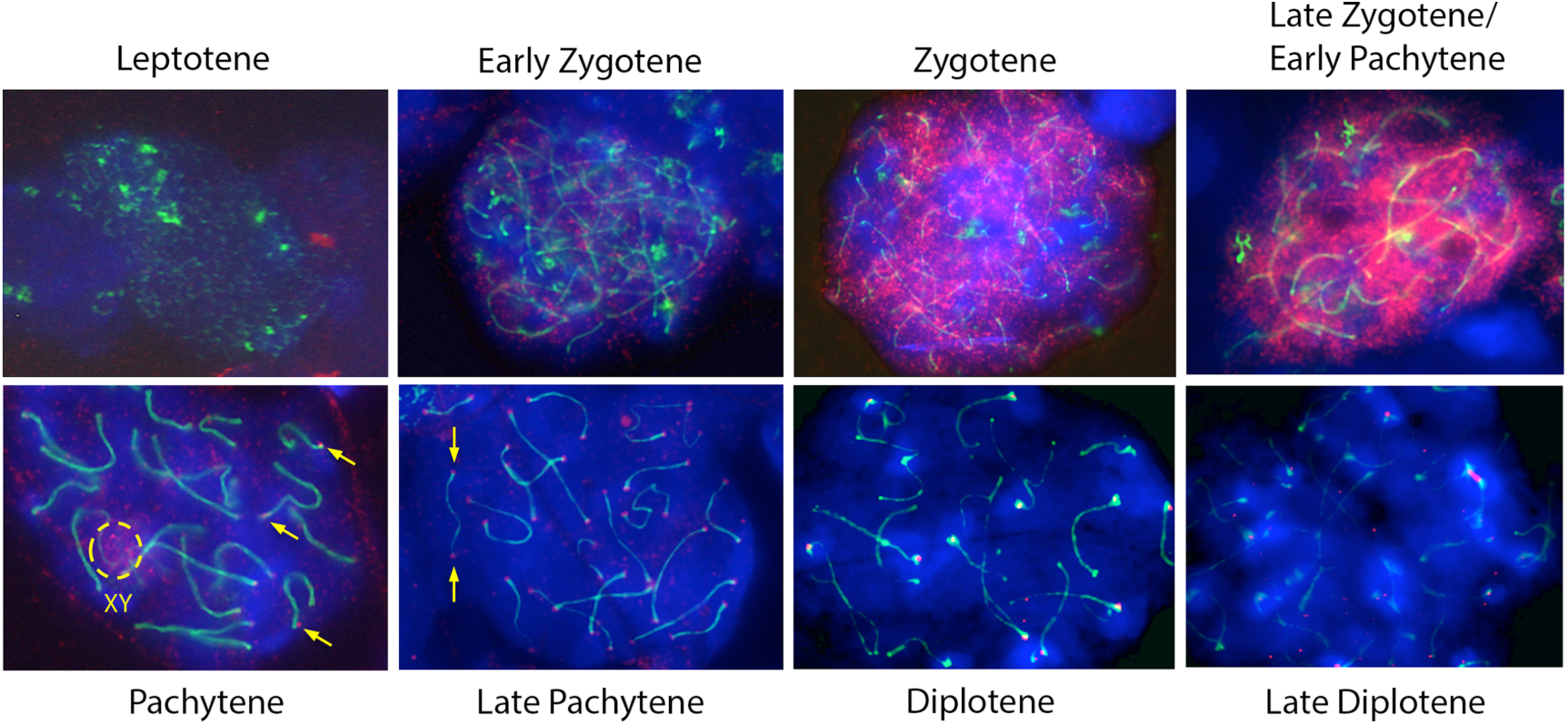
Expression of ZCWPW1 across meiosis prophase I in mouse testis. Nuclear spreads were immunostained with antibodies against ZCWPW1 (red) and the synaptonemal complex protein SYCP3 (green) which labels the chromosome axis, and counterstained with DAPI (blue) to visualise nuclei. Developmental stages indicated above. Yellow arrows show apparent ZCWPW1 foci at both ends of the synaptonemal complex. The dashed circle shows staining in the XY body.

We next studied mice from a constitutive *Zcwpw1* KO line (Methods), carrying a ∼1.5kb frameshift deletion encompassing exons 5 to 7 upstream of the CW domain, creating a premature stop codon resulting in the production and predicted degradation (by RNA-mediated decay) of a short 492bp (vs 1893bp for WT) transcript (**Figure 3A**). Confirming this, ZCWPW1 expression was completely absent in testis chromosome spreads from *Zcwpw1^−/−^* mice, in zygotene cells (**Figure 3B**) and all other meiotic stages of prophase I where expression is detected in WT mice (**Supplementary Figure 4**). Confirming recent findings in a different *Zcwpw1^−/−^* mouse (M. Li et al., 2019), we observed no overt fertility phenotype in either sex in the heterozygous *Zcwpw1^+/−^* mice (data not shown). However *Zcwpw1^−/−^* male mice were sterile with complete azoospermia and reduced testis size (**Figure 3B,D**), while female mice retained fertility until around 7-8 months of age (**Supplementary Table 2**), and otherwise both sexes develop normally. As in (M. Li et al., 2019), we observe widespread asynapsed chromosomes in male mice, marked by γ-H2AX and HORMAD2, persistent DMC1 foci marking unrepaired DSBs (**Figure 4**), no meiotic progression beyond (pseudo)pachytene, failure to form the sex body, and a complete absence of MLH1 foci marking recombination crossover sites (**Supplementary Figure 5**).

**Figure 3.**
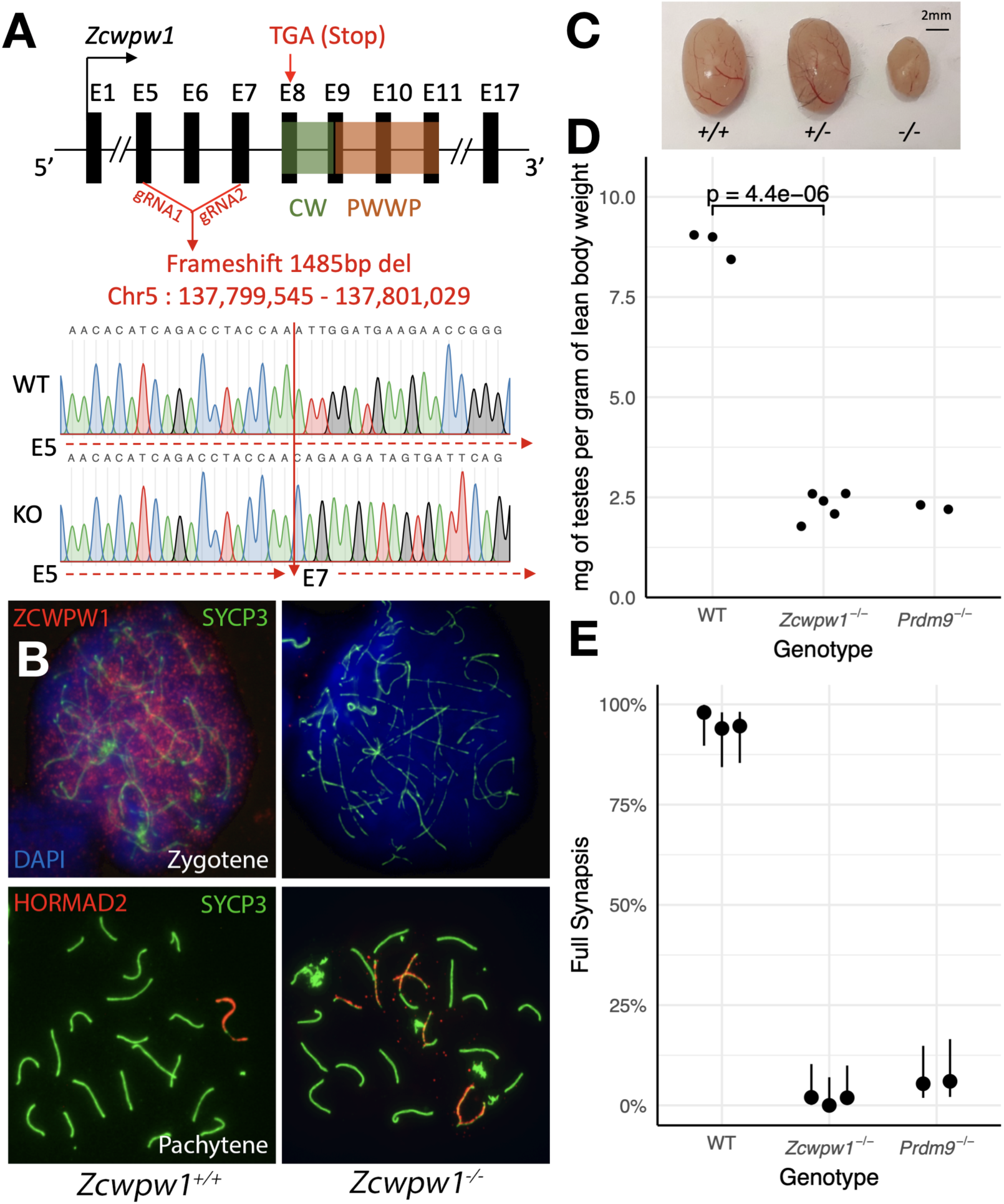
**Male *Zcwpw1^−/−^* mice show reduced testis size and asynapsis, similar to the *Prdm9^−/−^* mutant. (A)** Schematic of the *Zcwpw1* knockout mouse line. E: Exon. gRNA guideRNA. Sanger sequencing DNA chromatograms of wild-type (WT) and knockout (KO) mice encompassing the deletion are shown. The intron-exon organization is not to scale. **(B)** Immunofluorescence staining of testis nuclear spreads from *Zcwpw1^+/+^* and *Zcwpw1^−/−^* mice for ZCWPW1, the synaptonemal complex protein SYCP3 which labels the chromosome axis, or HORMAD2 which marks unsynapsed chromosomes. **(C)** Representative testes from 9-10 weeks old WT (+/+), Het (+/−) and Hom (−/−) *Zcwpw1* KO mice are shown. **(D)** Paired testes weight was normalized to lean body weight. The p-value is from Welch’s two-sided, two sample t-test. Raw data in **Supplementary Table 3**. **(E)** Synapsis quantification in testis chromosome spreads immunostained with HORMAD2, as in (B). The percentage of pachytene cells with all autosomes fully synapsed is plotted by genotype; n≥50 cells. Vertical lines are 95% Wilson binomial confidence intervals. Raw data in **Supplementary Table 4**.

**Figure 4.**
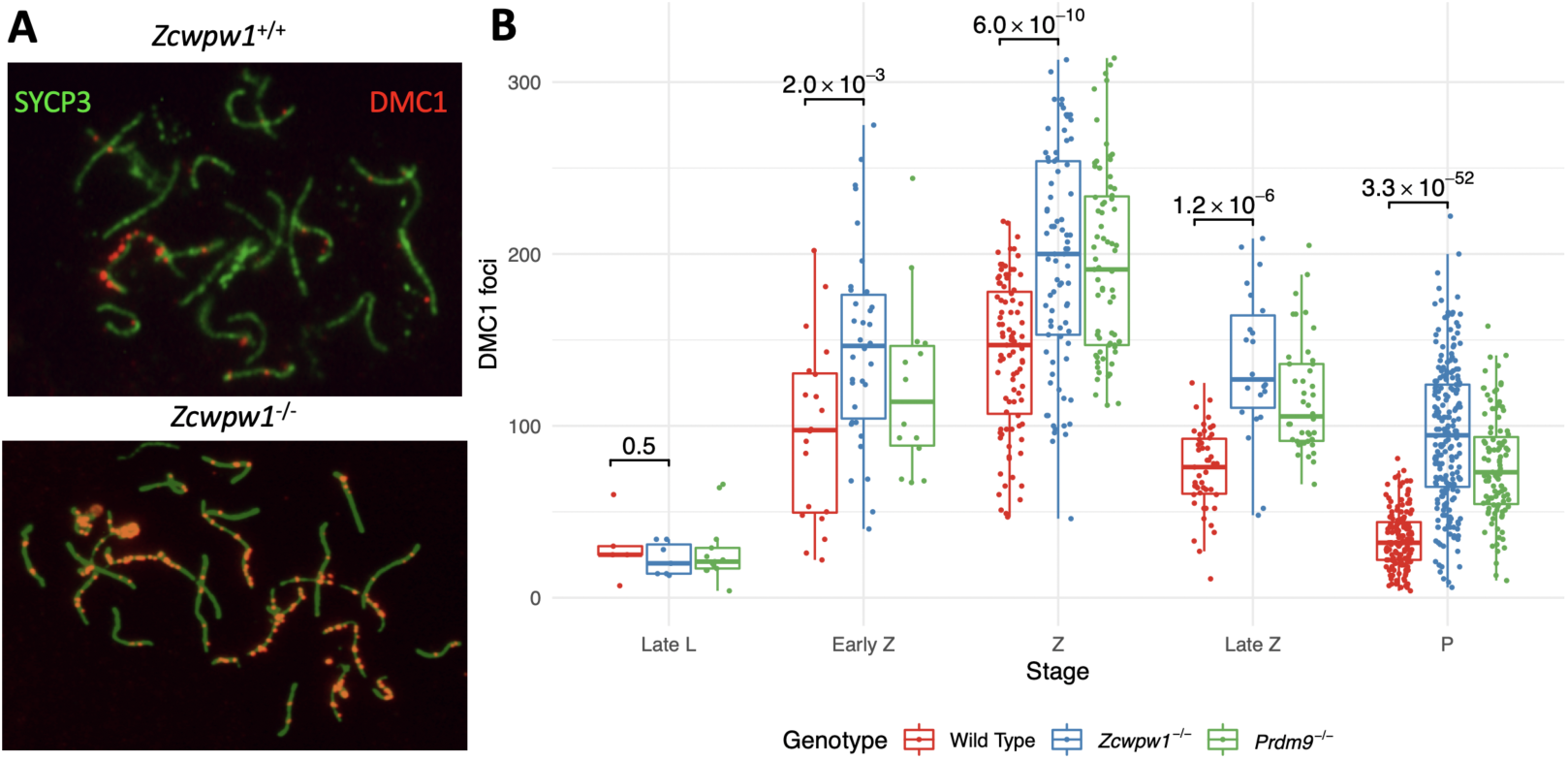
**Similar DMC1 count elevation in *Zcwpw1^−/−^* and *Prdm9^−/−^* mice, compared to wild-type. (A)** Testis chromosome spreads from wild-type and *Zcwpw1^−/−^* mice were immunostained for DMC1 and SYCP3. Representative late pachytene cells are shown. **(B)** The number of DMC1 foci in cells from the various stages of prophase I were counted. p-values are from Welch’s two sided, two sample t-test. L: Leptotene, Z: Zygotene, P: Pachytene. n=3 mice per genotype (*Zcwpw1^−/−^* and WT), n=2 for Prdm9^−/−^.

Each of these properties resembles observations in the *Prdm9^−/−^* mutant, and so we compared to this mutant. In our *Zcwpw1^−/−^* male mice, >98% of pachytene cells failed to properly synapse at least one pair of chromosomes (**Figure 3B, E**), similar to *Prdm9^−/−^* males. However, the nature of the synaptic defects observed differed (**Supplementary Table 4**). In the *Prdm9^−/−^* mutant, 63.2% of the pseudopachytene cells showed mispairing of non-homologous chromosomes in a typical branched structure (referred to as “tangled”, **Supplementary Table 4**). In contrast, we only observed 24.8% of *Zcwpw1^−/−^* pseudopachytene cells with this type of error, while the majority (74.5%) of cells contained multiple bundles of HORMAD2-positive unsynapsed chromosomes, which resemble the XY body and may merge with the sex chromosomes (thus referred to as “multibodies”, **Supplementary Figure 5**). These results imply that the *Prdm9^−/−^* mutant often mispairs chromosomes, while *Zcwpw1^−/−^* spermatocytes mainly fail to pair a subset of chromosomes at all. The expression levels and staining pattern of ZCWPW1 were not visibly altered in testis nuclear spreads from *Prdm9^−/−^* mice (data not shown).

Comparing levels of DMC1 foci as a proxy for DSB repair, or a potential cause for the asynapsis in *Zcwpw1^−/−^* males, in both *Zcwpw1^−/−^* and *Prdm9^−/−^* mice foci count was significantly elevated from early zygotene onwards, indicating delayed repair of DSBs (**Figure 4**). However, we observed a wider spread for *Zcwpw1^−/−^* males. A similar increase was observed at pachytene stage in the levels of RAD51 (**Supplementary Figure 6**), another ssDNA-binding protein which functions in concert with DMC1, with the large majority of RAD51 and DMC1 foci co-localizing (Brown et al., 2015; Tarsounas et al., 1999). In contrast, there was no difference in the levels of RPA2 (**Supplementary Figure 7**). In *Zcwpw1^−/−^* males, like in the *Prdm9^−/−^* mutant, DSBs form and recruit RPA2, RAD51 and DMC1 in similar numbers (M. Li et al., 2019), but fail to repair efficiently, accompanied by asynapsis and meiotic arrest at pseudo-pachytene. Indeed, we observe late unrepaired DMC1 foci mainly on asynapsed chromosomes (**Supplementary Figure 8**).

We also performed analyses of which chromosomes fail to synapse. FISH analysis revealed that chromosomes 18 and 19 are most often among these asynapsed chromosomes, while grouping of chromosomes based on their size revealed some asynapsis of all chromosomes, but at higher levels for shorter chromosomes (**Supplementary Figure 9**). This is similar to results observed for infertile hybrid male mice (Bhattacharyya et al., 2013), whose asynapsis is driven by lack of binding to the homologous chromosome by PRDM9 (Davies et al., 2016).

### ZCWPW1 is recruited to PRDM9 binding sites in an allele-specific manner

We previously studied the binding properties of human PRDM9 and established a genome-wide map in transfected human mitotic (HEK293T) cells by ChIP-seq (Altemose et al., 2017), observing binding to the majority of human meiotic recombination hotspots. Based on the presence of H3K4me3 and H3K36me3 recognition domains in ZCWPW1, we hypothesized that it would be recruited to PRDM9-bound genomic sites, where these marks are deposited upon binding in HEK293T cells (Altemose et al., 2017).

To test this, we co-transfected HEK293T cells with full-length human HA-tagged ZCWPW1 and either no other protein, or full-length PRDM9 alleles carrying the human or chimpanzee ZF-array, as studied previously (Altemose et al., 2017), and then performed ChIP-seq against the ZCWPW1 tag. Confirming the recruitment hypothesis, in the presence of human PRDM9 and compared to cells lacking human PRDM9, ZCWPW1 shows a strong enrichment at human PRDM9 binding sites (**Figure 5A, B**). Notably, even without PRDM9 we observed sequence-specific binding of ZCWPW1 to many sites in the genome (**Figure 5C and Supplementary Figure 10**). However, upon transfection, 91% (98%) of the strongest 10,132 (3,016) peaks are PRDM9 binding sites (cf. 7% expected overlap with randomised peaks), so PRDM9 is able to strongly reprogram ZCWPW1 binding, suggesting that non-PRDM9 peaks are bound more weakly. Because our transfection has <100% efficiency (**Supplementary Figure 11**), some cells containing ZCWPW1 will not possess PRDM9, and so it is possible that an even greater fraction of ZCWPW1 is redirected to PRDM9 binding sites in cells where both proteins *are* present.

**Figure 5.**
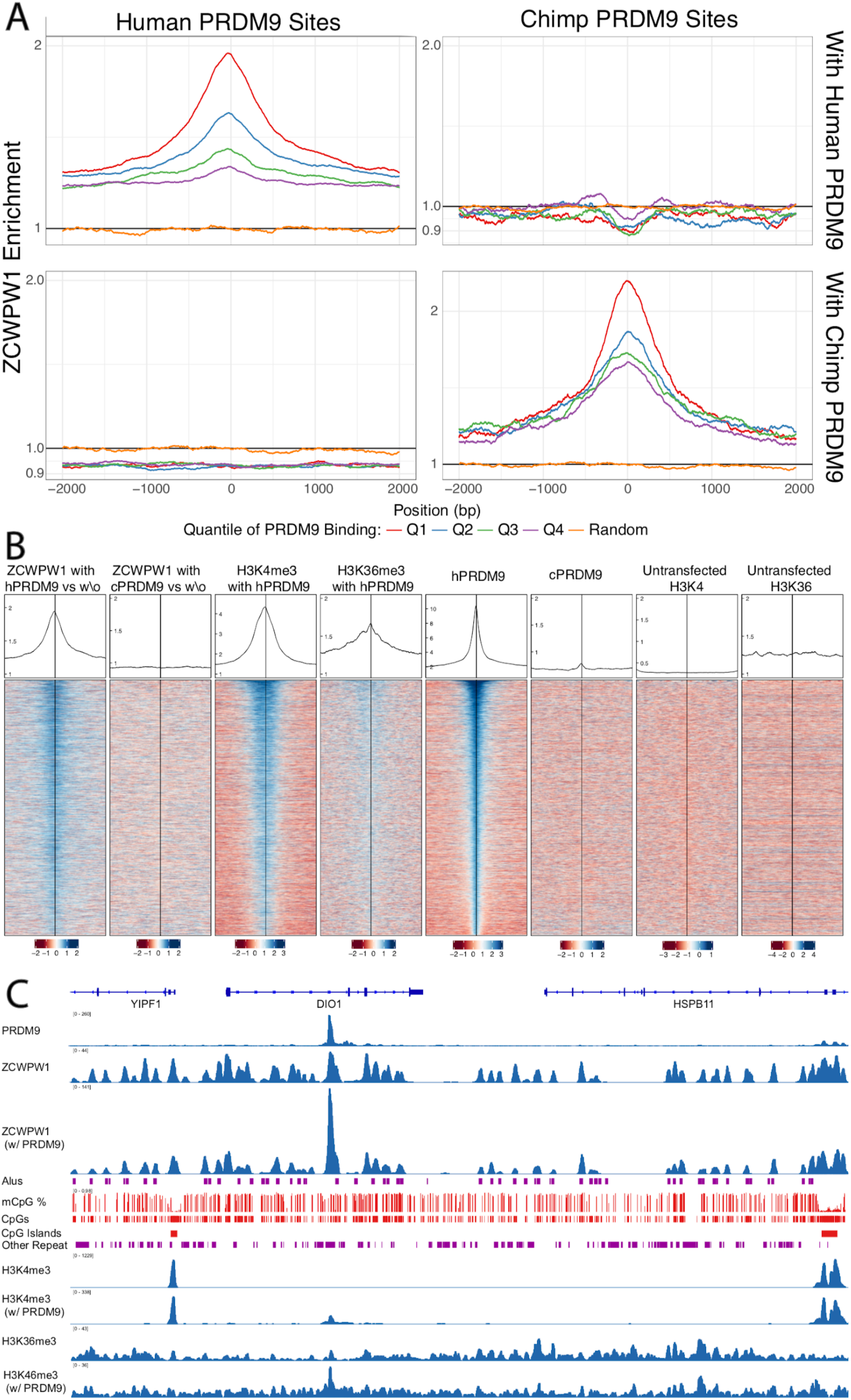
(**A**) Enrichment of ZCWPW1 (with vs without PRDM9) at PRDM9 binding sites when co-transfected with PRDM9 with either Human or Chimp Zinc Finger. Q = quartile. Human PRDM9 sites are centered and stranded by the motif. (**B**) Profiles and heatmaps of reads around top 25% of individual human (h) PRDM9 binding sites (rows). Heatmaps: log-fold change of target (indicated in column titles, Methods) vs input, for various labeled target proteins, ordered by human PRDM9. ZCWPW1, H3K4me3 and H3K36me3 each become enriched at human PRDM9 sites, following (co-)transfection with human PRDM9. (**C**) ChIP-seq data and annotation in a genome plot illustrate the behaviour of ZCWPW1 and other factors. ChIP-seq tracks show fragment coverage. Tracks where PRDM9 is present are labelled “w/ PRDM9”, and below corresponding tracks with out PRDM9. ZCWPW1 binds to Alus, CpG islands and other CpG-rich sequences even in the absence of PRDM9. On addition of PRDM9, ZCWPW1 becomes strongly enriched at PRDM9 binding locations (center left peak within DIO1).

Importantly, when co-transfecting with a modified version of PRDM9 in which the zinc finger array is replaced with that from chimp (which binds different locations in the genome (Altemose et al., 2017)), we find that the enrichment at human binding sites disappears, but instead ZCWPW1 is enriched at *chimp* PRDM9 binding sites (**Figure 5A and Supplementary Figure 12**). This perturbation experiment provides strong evidence that PRDM9 *causes* recruitment of ZCWPW1 (as opposed to for example independent recruitment of both proteins). Notably, the strength of binding of ZCWPW1 at these sites provides a better predictor of DSB formation (DMC1 sites) than does PRDM9 binding strength itself (**Supplementary Figure 13**): although we show (see below) that ZCWPW1 is *not* directly involved in DSB positioning, this might suggest involvement of similar features to those it recognizes, in recruiting DSBs.

We tested whether this recruitment might be mediated by the dual histone modifications H3K4me3 and H3K36me3. Consistent with this idea, ZCWPW1 binding is positively associated with levels of both H3K4me3 and H3K36me3 marks (**Supplementary Figures 14 and 15**). We examined transcription start sites, which possess H3K4me3 at high levels, but lack H3K36me3, observing some ZCWPW1 signal at these sites, but with a uniformly lower mean ZCWPW1 enrichment compared to those with evidence of PRDM9 binding (and hence both histone marks) (**Figure 6 and Supplementary Figure 16**). Although it is not possible to measure H3K4me3 presence/absence in *individual* cells in our system, even the most weakly bound PRDM9 sites – which must therefore possess H3K4me3 in only a fraction of cells – show stronger ZCWPW1 enrichment than the strongest promoters, which are likely to have near 100% H3K4me3 marking, and possess >2-fold more H3K4me3 than even the strongest PRDM9 binding sites. We conclude that H3K4me3 alone endows only relatively weak binding, while PRDM9 is therefore able to recruit ZCWPW1 with a much greater efficiency than sites marked by H3K4me3 alone, suggesting that both histone modifications might aid efficient binding.

**Figure 6.**
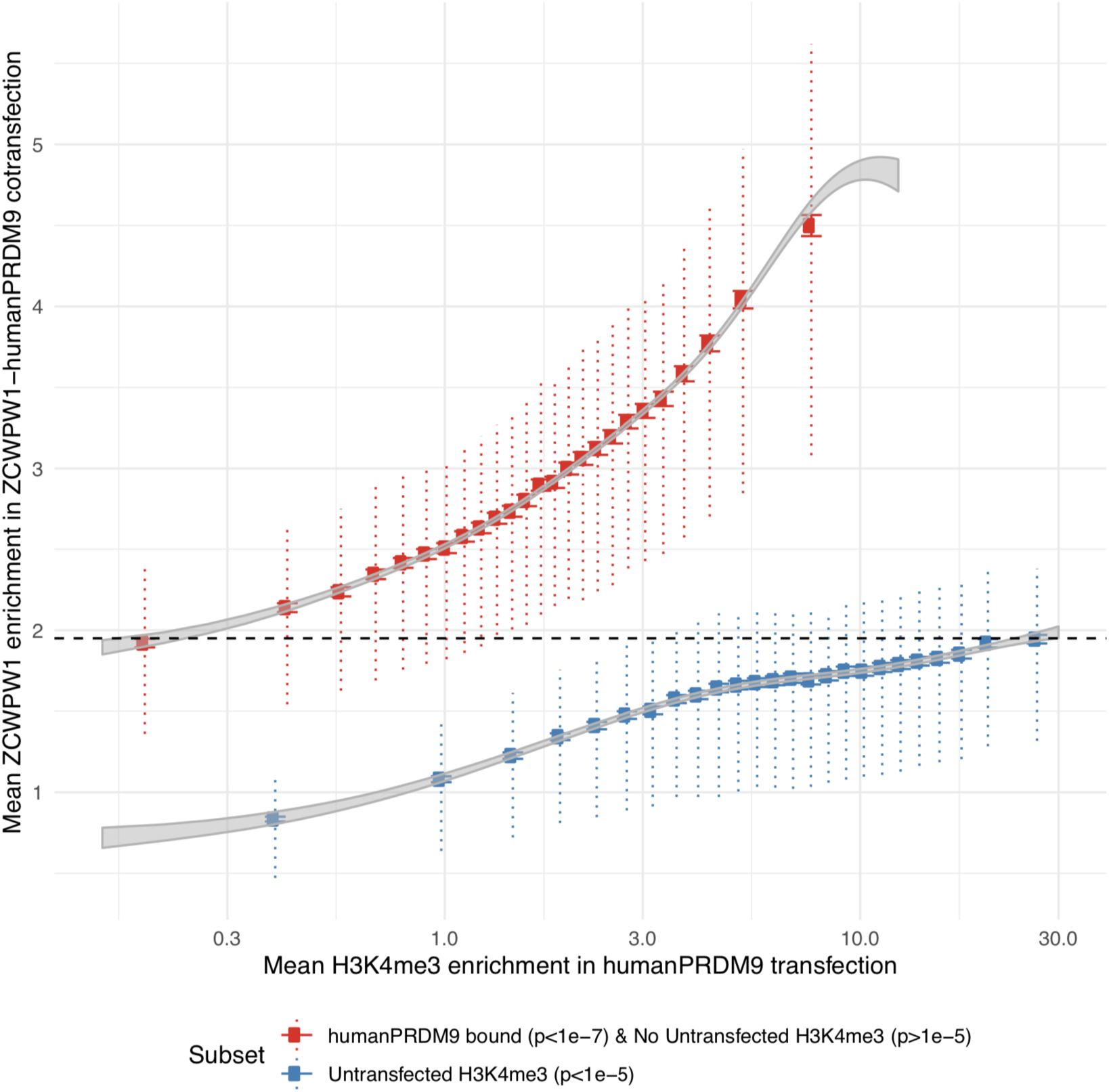
**PRDM9 (H3K4me3 and H3K36me3) is a stronger recruiter of ZCWPW1 than promoters (H3K4me3 only)**. H3K4me3 and ZCWPW1 were force called in 100bp windows. Each subset is defined as indicated in the legend with the additional constraint of requiring input fragment coverage >5 for ZCWPW1 and >15 for H3K4me3. For each subset H3K4me3 was split into 25 bins with equal number of data points. Horizontal bars: 2 standard errors of the mean. Vertical dotted bars: upper and lower quartiles. Grey ribbons show 2 standard errors for a Generalized additive model on log(mean H3K4me3 enrichment + 0.1). Dashed black horizontal line highlights that the mean enrichment of the highest bin for promoters is similar to that of the lowest bin for PRDM9 bound sites.

### DSBs occur at their normal locations in *Zcwpw1*^−/−^ mice but show altered DMC1 persistence

Previous work has shown that *Prdm9^−/−^* mice use a new set of DSB hotspots, localising at CpG islands and/or promoter regions (Brick et al., 2012). Given that PRDM9 recruits ZCWPW1, one possible function of ZCWPW1 may be that, in turn, it recruits the DSB machinery and hence forms part of the causal chain in normal positioning of DSBs at PRDM9-specified hotspots. Alternatively, the Z*cwpw1*^−/−^ mutant phenotypes we observe might reflect a more downstream role. To distinguish these hypotheses, we gathered data on DSB positioning and repair dynamics genome-wide, by carrying out single stranded DNA (ssDNA) sequencing (SSDS) by ChIP-seq against DMC1 (Khil et al., 2012).

In the Z*cwpw1*^−/−^ mutant, we observed normal localisation of DMC1 at the expected B6 hotspots for this background, with no activity at *Prdm9^−/−^* hotspots (**Figure 7A**). Moreover, individual wild-type hotspots appear always to be active in the Z*cwpw1*^−/−^ mutant (**Figure 7C**): we only fail to see evidence of DMC1 signal in a subset of the weakest hotspots, where we are likely to lack statistical power (**Supplementary Figure 17**). Thus DSBs occur in unchanged hotspot regions in Z*cwpw1*^−/−^ males. To check if they occur at the same locations *within* hotspot regions, we leveraged data for SPO11-mapped DSB sites (Lange et al., 2016). Specifically, we identified three sets of hotspots whose respective mapped breaks occur mainly upstream, central, or downstream of the PRDM9 binding site (Methods), and compared their DMC1 ChIP-Seq signal profiles, as well as their SPO11 signals (**Supplementary Figure 18**). This revealed that in the Z*cwpw1*^−/−^ males, as in the wild-type, DMC1 signals, though broader, mirror this break positioning. Therefore, ZCWPW1 is not required to specify hotspot locations, and neither does it strongly influence where DSBs occur *within* hotspots.

**Figure 7.**
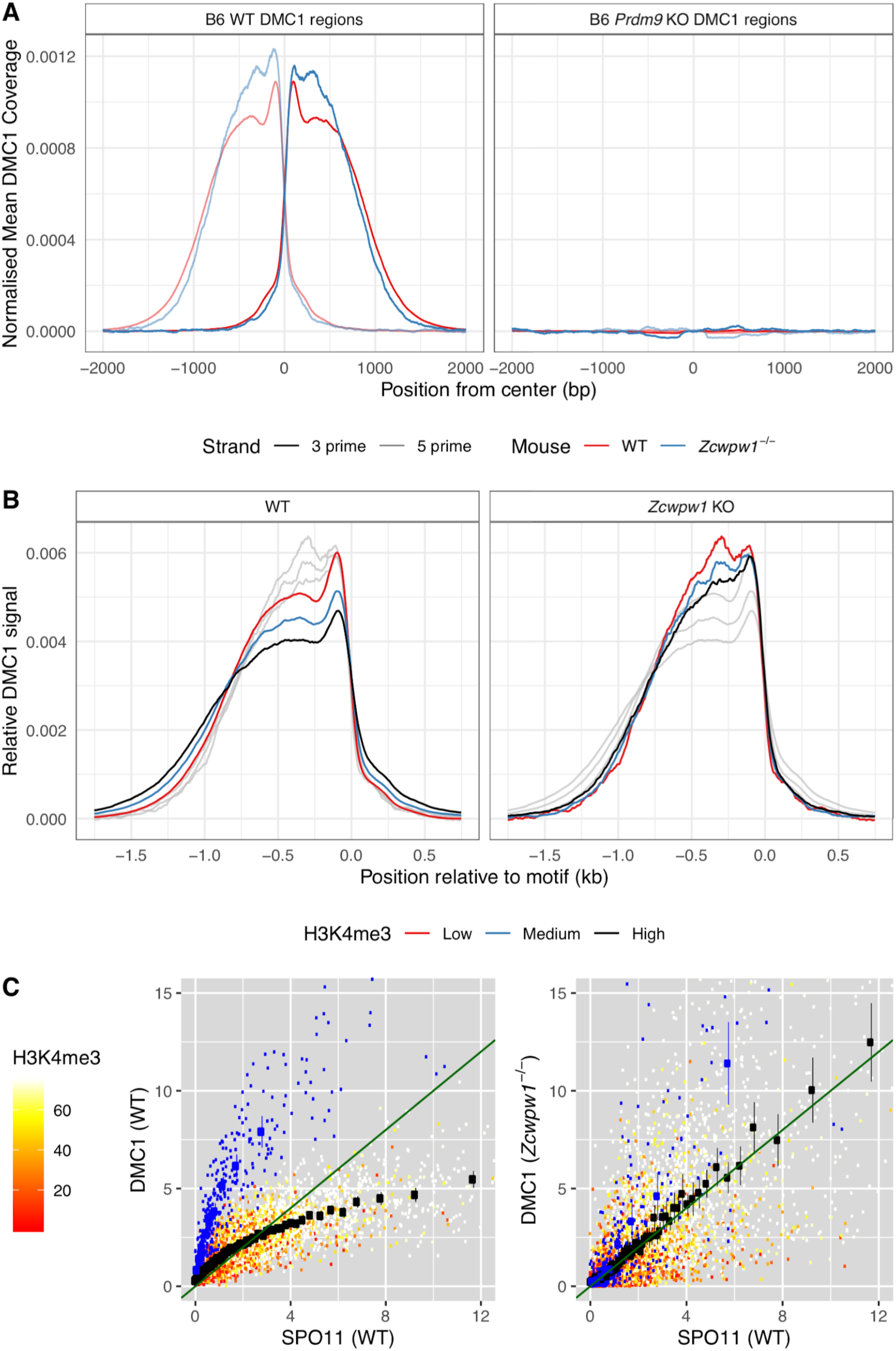
**(A)** DSBs occur at normal hotspot locations in the *Zcwpw1^−/−^* mouse. Average coverage of reads from DMC1 SSDS ChIP-seq at previously mapped regions (Methods) in B6 WT (left) and *Prdm9^−/−^* (right) mice is shown, centered at the PRDM9 motif (left). DMC1 profiles from a WT mouse are shown in red, data from (Brick et al., 2012). (**B**) Normalised DMC1 profile is plotted for WT and *Zcwpw1*^−/−^, stratified by H3K4me3. Greyed out lines show the alternative genotype for comparison. **(C)** Altered DMC1 signals in *Zcwpw1^−/−^* mice. Relationship between SPO11-oligos (measuring the number of DSBs) vs DMC1 (a measure of the number and persistence of DSBs) at each B6 hotspot for WT and *Zcwpw1^−/−^*. Green line is y=x for reference.

One perhaps important difference between the *Zcwpw1^−/−^* and wild-type DMC1 profiles is that the latter signal grows slightly (up to ∼200bp) wider in hotspots with increasing PRDM9-induced H3K4me3 (**Figure 7B**), while this effect is absent in the *Zcwpw1^−/−^* case. This might be explained by small differences in chromatin accessibility subtly impacting SPO11 locations, DNA end resection which generates the 3’ ssDNA tail to which DMC1 binds, or downstream processing differences altering DMC1 span in the mutant mice.

Despite very similar DSB locations, we saw much greater, and systematic, differences in the strength of the DMC1 signal at individual hotspots. As previously (Davies et al., 2016; Khil et al., 2012), we note that observed average DMC1 signal strength at a hotspot reflects the product of the frequency at which DSBs occur there, and the average length of time DMC1 remains bound to the ssDNA repair intermediate, with the latter reflecting DSB processing/repair time. In contrast, available SPO11 ChIP-Seq data (Lange et al., 2016) reflect mainly the former, the frequency of DSBs. We therefore compared DMC1 and SPO11 signal strength at each autosomal and X-chromosome hotspot, in WT and *Zcwpw1^−/−^* mice. In WT mice (as seen previously in other mice (Davies et al., 2016) including, albeit somewhat more weakly, even sterile hybrids) the non-PAR X-chromosome shows a very strong elevation of DMC1 signal strength, reflecting the persistent DMC1 foci on this chromosome also visible using microscopy, at DSB sites that eventually repair using the sister chromatid. Moreover, in WT mice hotspot heat (with hotter hotspots having stronger H3K4me3 signal also) shows a sub-linear relationship between SPO11 and DMC1. This is thought to reflect a wider phenomenon of faster DSB repair occurring within those hotspots whose homologue is more strongly bound by PRDM9, i.e. those hotspots with a stronger H3K4me3 signal in the WT mouse (Davies et al., 2016; Hinch et al., 2019; R. Li et al., 2019). The X-chromosome DMC1 elevation is in a sense an extreme case of slower repair of DSBs whose homologue is not PRDM9-bound, because no homologue exists in this case.

However, we see a striking departure in the *Zcwpw1^−/−^* mouse, where the X-chromosome behaves similarly to the autosomes in terms of DMC1 vs. SPO11 signal strength (**Figure 7C**). Moreover, we also see a simple linear relationship between DMC1 and SPO11 binding in this mouse. These data imply the relationship between DMC1 removal and PRDM9 binding is eliminated in this mouse, so PRDM9 (whether because ZCWPW1 directly aids the process, or indirectly) appears unable to “assist” in homology search in this mouse. Indeed, the data imply a widespread perturbation of DSB repair in this mouse, with autosomal DMC1 foci persisting as long as those on the X-chromosome. In the previously studied *Hop2*^−/−^ mouse (Khil et al., 2012; Petukhova et al., 2003; Smagulova et al., 2011), we identified a very similar pattern (**Supplementary Figure 19**): in this mouse, DMC1 is loaded onto ssDNA, but DSBs are not repaired at all, indicating that DSB repair pathways in *Zcwpw1^−/−^* mice are profoundly altered. These results are consistent with, and extend, our microscopy observations (**Figure 4**) that many DSBs persist in the *Zcwpw1^−/−^* mouse, and some may never repair. We are unable to say whether DSB repair involves the homologue or not in this mouse, but the partial synapsis we observe suggests some repair does likely occur. To further understand DMC1 persistence at individual hotspots, we estimated the relative DMC1 heat of 9,318 autosomal hotspots in the *Zcwpw1^−/−^* mouse, compared to the WT mouse (Methods). Via linear regression, by far the best individual predictor of this ratio among H3K4me3, WT DMC1, and SPO11 was the level of H3K4me3 (r=0.55, p<10^−15^; **Supplementary Figure 20**, while r=0.57 when using all 3 predictors together). The average DMC1 ratio changes around 9-fold from the most weakly to the most strongly H3K4me3-marked hotspots. Therefore, if DMC1 signal changes are indeed explained by slower DSB processing in the *Zcwpw1^−/−^* mouse at hotspots bound strongly by PRDM9, then this implies a very strong effect. Whatever the cause, the overall strong correlation implies perturbation of DMC1 behaviour is widespread, rather than impacting any small subset of hotspots. Moreover, it appears to be mainly controlled by local levels of PRDM9 histone modification, in keeping with the evidence that ZCWPW1 recognizes this mark at PRDM9-bound sites, on either the broken or (identical) homologous chromosome.

### ZCWPW1 binds CpG dinucleotides

In addition to the strong PRDM9-dependent ZCWPW1 peaks described earlier, there are many locations in HEK293T cells at which ZCWPW1 binds, typically more weakly, and independently of PRDM9. Indeed, we identified over 800,000 ZCWPW1 peaks. Surprisingly, a large proportion of these binding sites overlap Alu repeats (**Figure 8A and Supplementary Figure 21**) (of which there are 1.1 million in the human genome (Deininger, 2011)). The weakest ZCWPW1 peaks overlap Alus most frequently, whilst the strongest peaks are depleted of Alus relative to chance overlap (**Figure 8A**).

**Figure 8.**
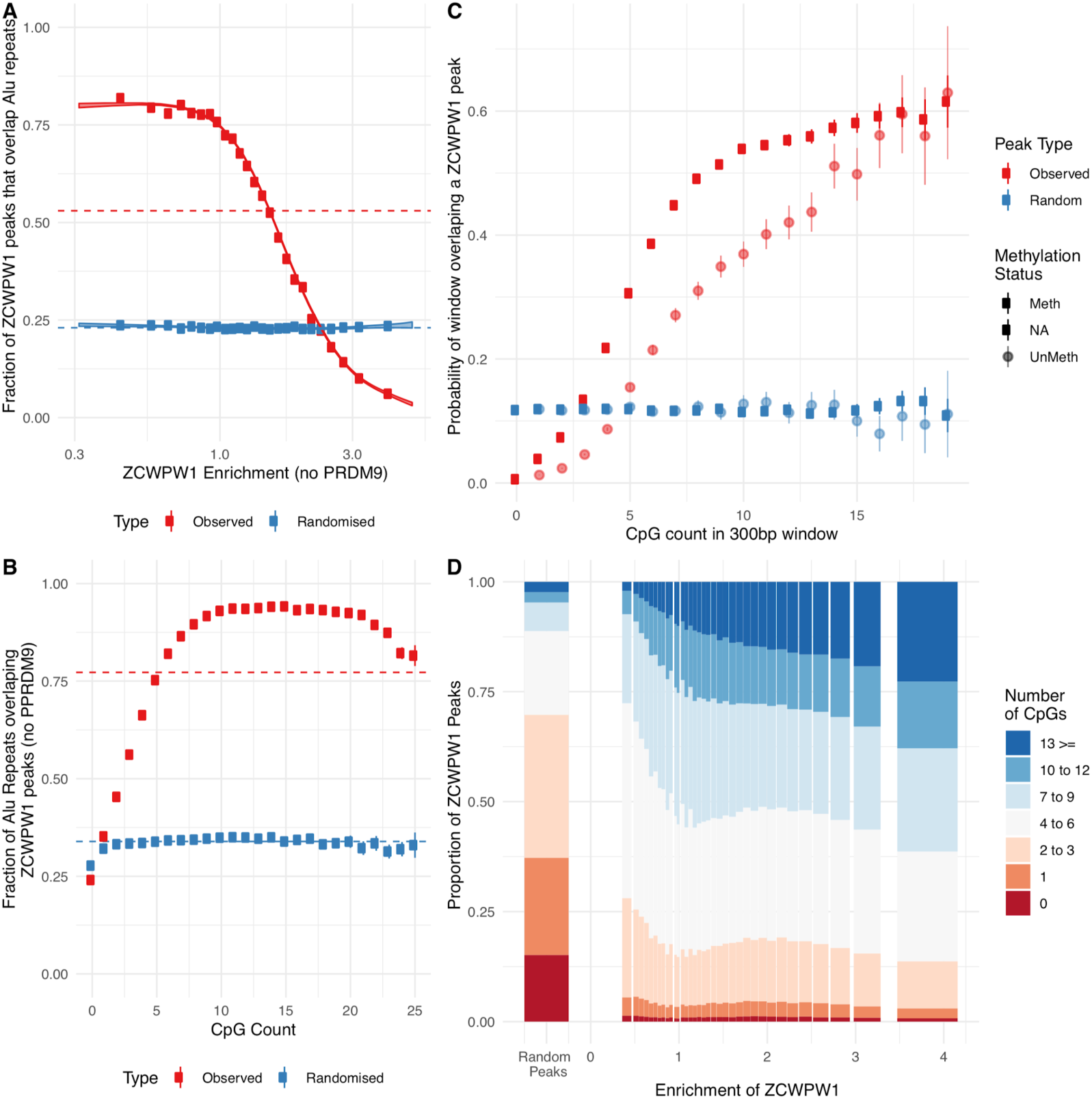
ZCWPW1 binds CpG-rich sequences such as Alu repeats. **(A)** Fraction of overlap of ZCWPW1 binding peaks with Alus, ordered by enrichment in ZCWPW1 binding. ZCWPW1 peaks are binned into 25 bins with equal number of data points, and means of both enrichment and overlap are plotted. Solid ribbon shows prediction from GAM logistic model. Dotted line shows overall means. **(B)** Rate of overlap of Alu repeats with ZCWPW1 peaks, for Alus with different numbers of CpG dinucleotides. **(C)** The probability of a 300bp window overlapping a ZCWPW1 peak increases with increasing CpG count in that window. Windows overlapping (by 10bp or more) Alus, other repeats, or CpG islands have been excluded. Methylation status of NA is for windows with 0 CpGs. Methylated regions are those with a methylated to unmethylated reads ratio of >0.75, and unmethylated <0.25 (Methods) **(D)** Relative proportion of peaks with given numbers of CpGs +/−150bp from peak center. ZCWPW1 peaks are enriched in CpGs compared to random peak locations. ‘Meth’, methylated; ‘UnMeth’, unmethylated; NA, not applicable.

Because Alus are rich in CpGs (containing >23% of human CpG dinucleotides, with deamination depleting CpGs in older Alus (Luo et al., 2014)), we tested if binding correlates with CpG presence/absence. Indeed, the binding of ZCWPW1 to Alus depends on the presence of CpGs, which are mainly methylated (Gaysinskaya et al., 2018): Alus with no CpGs were bound at a rate lower than by chance, but almost all Alus containing 10-20 CpGs are bound (**Figure 8B**). This dependence is not Alu-specific: even outside genomic repeats, 300-bp regions with zero CpGs have negligible probability of overlapping a ZCWPW1 binding site, while the overlap probability rises to >50% for regions containing 10 or more CpGs, suggesting CpGs are essential for PRDM9-independent ZCWPW1 binding (**Figure 8C,D and Supplementary Figure 22**). ZCWPW1 appears to have greater affinity for methylated CpG pairs, but retains some affinity even for non-methylated regions. This binding mode was unexpected *a priori*, and does not appear to be easily explained by patterns of H3K4me3/H3K36me3 in the genome, which do not concentrate as strongly in these regions.

## Discussion

Here we exploited previous work on single-cell transcript profiling of mouse testis, which has established spatio-temporal (and likely functional) components of co-expressed genes (Jung et al., 2019), to identify co-factors relevant to PRDM9 function in specification of recombination hotspots and DSB repair during meiosis. We show that the histone methylation reader ZCWPW1, which is both highly co-expressed and co-evolving with PRDM9, is recruited by PRDM9 to its binding sites, likely mediated through the recognition of the dual H3K4me3-H3K36me3 histone mark PRDM9 deposits.

While we find co-evolution of ZCWPW1 with PRDM9, consistent with proteins functioning in the same pathway, this appears to depend on particular domains. ZCWPW1 shows weaker conservation among species outside the CW and PWWP domains, suggesting these are of high importance, and co-occurs with PRDM9 in species where the latter possesses a SET domain predicted to retain catalytic activity for H3K4me3/H3K36me3, which the domains of ZCWPW1 are predicted to recognise, strongly suggesting a partnership totally dependent on this recognition. This co-occurrence can persist, even in species where PRDM9 is not believed to position DSBs, suggesting the co-operation between these proteins does not completely depend on this aspect of PRDM9’s function. Interestingly, such a SET domain of PRDM9 mainly occurs when PRDM9 possesses a SSXRD domain, which in other proteins (Banito et al., 2018) recruits CXXC2 which binds to CpG islands – whether this connects to our observation of a potential CpG-binding domain in ZCWPW1, and CpG-binding behaviour *ex-vivo*, remains to be explored.

Others have recently reported phenotyping of *Zcwpw1^−/−^* mice (M. Li et al., 2019), and although our KO is different, we observe similar patterns of infertility in males, and results consistent with gradual fertility decline in females, towards sterility from 8 months onwards. Why females deficient for ZCWPW1 are (at least initially) fertile is unknown, but resembles sexual dimorphism in loss-of-function mutants for other meiotic genes involved in homologous recombination, where females exhibit a milder fertility phenotype (Cahoon and Libuda, 2019; Morelli and Cohen, 2005; Zhang et al., 2019). However we observe a distinct intracellular localisation of ZCWPW1 from that reported by (M. Li et al., 2019): while we also detect ZCWPW1 as a diffuse nuclear signal across the nucleus (though excluding the chromocenters) and localisation to the XY body in pachytene, we see an additional signal in mid-pachytene to diplotene cells at the ends of the synaptonemal complex. This might reflect the specificity of our polyclonal antibody, which is raised against the full-length mouse protein vs a monoclonal antibody raised against a C-terminal region, hence recognising a wider spectrum of ZCWPW1. Given that the *Zcwpw1^−/−^* mouse arrests in early pachytene, it is not possible to test later stages, so a caveat is that we cannot completely exclude off-target antibody recognition. If accurate, the late-stage localisation of ZCWPW1 at subtelomeric regions resembles other proteins (Speedy A, CDK2, TFR1, SUN1, TERB1/2, MAJIN, and KASH), which form a complex tethering the telomeres to the nuclear envelope to facilitate chromosome movement and pairing - essential for synapsis and fertility (Ashley et al., 2001; Ding et al., 2007; Horn et al., 2013; Shibuya et al., 2014; Tu et al., 2017). However, no evidence of impaired telomeric attachment was reported in the other *Zcwpw1^−/−^*, based on normal localisation of TRF1 (M. Li et al., 2019).

We find, using an *ex vivo* system, that ZCWPW1 is strongly recruited to PRDM9-bound sites, in an allele-specific manner, dependent on the level of H3K4me3 and H3K36me3 that PRDM9 deposits, and most likely due to recognition of these marks by the CW and PWWP domains of ZCWPW1. Li and colleagues (M. Li et al., 2019) showed co-immunoprecipitation of H3K4me3 with ZCWPW1, but our results imply that PRDM9 recruits ZCWPW1 much more strongly than H3K4me3 alone, further suggesting that the *combination* of histone marks is key for ZCWPW1’s recognition of PRDM9 sites. Indeed in cells with PRDM9, the strongest ZCWPW1 binding sites are almost all PRDM9-bound sites.

Although it has been suggested that ZCWPW1 might recruit the DSB machinery (M. Li et al., 2019; Spruce et al., 2019), our data imply DSB formation and positioning are largely unaffected by loss of ZCWPW1 and occur at PRDM9-bound hotspots. Moreover DMC1, RAD51 and RPA2 loading also appear normal (**Figure 7A** and (M. Li et al., 2019)), but asynapsis and failure to remove DMC1 imply a profound downstream impact. Indeed most hotspots show perturbed DMC1 heats consistent with DSB repair delays proportional to their *cis* H3K4me3 levels, i.e. driven by expected WT ZCWPW1 recruitment levels at each hotspot (**Supplementary Figure 15**). ZCWPW1 therefore offers a (thus far) unique protein, linking PRDM9 binding with its downstream functions (Davies et al., 2016; Hinch et al., 2019; R. Li et al., 2019) in aiding rapid DSB processing/repair at hotspots whose homologue is bound by PRDM9.

PRDM9 is thought to aid homology search/chromosome pairing by such binding to the homologous chromosome, and hybrids where hotspots are highly “asymmetric” (i.e. the homologues are not bound at the sites where DSBs occur, due to evolutionary hotspot erosion) show multiple features identical to Zcwpw1^−/−^ mice. These include asynapsis (particularly of shorter chromosomes with fewer DSBs), persistent DMC1 foci (at the asymmetric hotspots), and complete sterility in males only. In Zcwpw1^−/−^ mice, we infer that the level of PRDM9 binding to the homologue now makes no difference to DSB persistence, so these mice generally behave exactly as if all hotspots are completely asymmetric. Thus, a parsimonious explanation of our findings is that ZCWPW1 functions by attaching to PRDM9-bound sites, so as to recognize the homologous chromosome at DSB sites. This might e.g. lead to their recruitment to the chromosomal axis (where DMC1 foci are found), for potential use as the homologous repair template in DSB repair, perhaps mediated by the SCP-1-like domain identified in some orthologous copies. Given we observe ZCWPW1 throughout the nucleus in early meiosis, such recruitment would likely be transient in nature. Alternatively, ZCWPW1’s part in mediating PRDM9’s downstream roles might also depend (or even, depend only) on it binding the chromosome on which DSBs occur.

Given that we still observe some synapsis, similar to *Prdm9^−/−^* mice, we suggest that a “back-up” mechanism, which is independent of PRDM9 binding level, is acting to attempt homology search/chromosome pairing in *Zcwpw1^−/−^* mice. Some such mechanism is necessary, to repair DSBs happening within a subset of hotspots that are asymmetric due to naturally occurring variation (as seen in e.g. humans and heterozygous mice). If our observed >9-fold changes in relative DMC1 levels for strongly vs. weakly bound hotspots purely reflect increased DMC1 persistence, it might be this mechanism is greatly slower than the PRDM9-dependent pathway for wild-type mice. In turn, this could explain many – or all – of the downstream phenotypes in these mice, as consequences of delayed homology search.

The simplest explanation of our observation that ZCWPW1 binds in a CpG dependent manner *ex vivo* is that these sites might be directly bound by a methyl-CpG binding domain (MBD) putatively observed in the human, mouse, and coelacanth *Zcwpw1*, among others. Alternatively, recruitment might be indirect, with ZCWPW1 forming a complex with an MBD containing protein, e.g. SETDB1 which is expressed in HEK293 cells, co-expressed with ZCWPW1 and PRDM9 (Jung et al., 2019; Schultz et al., 2002), and interacts with several other identified PRDM9 partner proteins (Mulligan et al., 2008; Parvanov et al., 2017). Given the precisely regulated developmental changes in methylation status, ZCWPW1/PRDM9 protein abundance, and chromatin during meiosis (Gaysinskaya et al., 2018; Seisenberger et al., 2013) it is unclear whether ZCWPW1 will also weakly bind Alus *in vivo*. However, if CpG dinucleotides do retain weak *in vivo* ZCWPW1 binding affinity, and given that CpG methylation is associated with transposons, it is interesting to speculate whether ZCWPW1 might play a role in germline transposon recognition, silencing, or facilitate repair of DSBs occurring within transposable elements.

*Zcwpw1* possesses a paralogue, *Zcwpw2*, which is also co-expressed in testis with *Prdm9* (although at a lower level), possesses both CW and PWWP domains, and has been shown to be able to recognise H3K4me3 at least (Liu et al., 2016). Further studies are required to investigate what function if any *Zcwpw2* has in meiosis. One interesting possibility is that *Zcwpw2*, rather than *Zcwpw1*, might act to help position DSBs – if so, its low abundance relative to *Zcwpw1* would be analogous to the low number of DSBs per meiosis (∼300) relative to the number of PRDM9 binding sites.

In this study we have shown that *Zcwpw1* co-evolves with *Prdm9*, binds to sites marked by PRDM9, and is required for the proper processing, removal of DMC1, and repair of DSBs. How exactly it mediates repair, if for example by recruitment of other proteins, remains to be determined. This study also further demonstrates that genes co-expressed with PRDM9 represent a rich source of undiscovered meiotic genes, with important functional implications for recombination, meiosis progression and ultimately fertility.

## Methods

### Orthologue Alignment

We identified ZCWPW1 orthologues across species using four data sources: first, we used BlastP (blast.ncbi.nlm.nih.gov), against the full-length human reference ZCWPW1 sequence (identifier NP_060454.3, against nr_v5 database) storing the top 1000 hits using the default parameters (set 1). Secondly, we downloaded the two sets of Ensembl identified ZCWPW1 orthologues (against ENSMUSG00000037108.13; ensembl.org, 105 orthologues), and identified NCBI ZCWPW1 protein orthologues (146 species, www.ncbi.nlm.nih.gov). Initial examination of identified orthologues revealed conservation among orthologues mainly of the ZW and PWWP domains; therefore, to find additional orthologues sequences we performed tBLASTn (against the nr/nt nucleotide collection 17^th^ July 2019; top 1000 hits) to identify orthologues in the NCBI nucleotide database, to the partial sequence (amino acids 256-339 of the reference sequence NP_060454.3), corresponding to the ZW and PWWP domains of ZCWPW1. Protein sequences were then aligned against full-length human ZCWPW1 (NP_060454.3) using BLASTP2.9 (Altschul et al., 2005, 1997) We obtained taxonomy information from the folder https://ftp.ncbi.nlm.nih.gov/pub/taxonomy/new_taxdump/, for comparisons among species and between ZCWPW1 and PRDM9.

Previous work has identified highly conserved “KWR” and “PWWP” patterns among ZW and PWWP domains, respectively (Fahu He et al., 2010; Qin and Min, 2014). We thus identified an initial set of “clear” ZCWPW1 orthologues, and then used these to identify further less precise matches. “Clear” orthologues are defined as proteins within set 1 containing perfect matches to both these sequences, and such that in the NCBI alignment, at least 39% of each sequence aligns to the human protein, and conversely. This second step is required to avoid spurious matches to e.g. ZCWPW2, which overlaps 19% of ZCWPW1, by requiring >2-fold this match length; although several ZCWPW2 copies are in our initial list, no gene annotated as most similar to ZCWPW2 attains the 39% overlap. For proteins within the “clear” set, we identified the longest alignment for each given species, resulting in 136 species with a likely ZCWPW1 orthologue. This initial screen identified three groups of fish, placental mammals, marsupials, monotremes and reptiles as likely possessing ZCWPW1, similar to our final conclusions. Moreover, it defined amino acids 247-428 in the human reference sequence as being conserved in the alignment (at most 1 sequence not aligning), including the annotated ZCW and PWWP domains.

Using this initial set, we identified additional orthologous sequences and refined our results, by identifying conserved bases within ZCWPW1. Specifically, we divided the 136 species into major clades, as in (Baker et al., 2017), and gave each sequence a weighting so that the overall weight for each clade was the same (so e.g. each Placental mammal sequence was downweighted as this clade was over-represented). Within the consistently aligned region 247-428, we calculated the overall weighted probability of each of the 20 amino acids, or a gap (adding 10^−5^ to exclude zero weights). This identified 31 completely conserved amino acids, and 78 amino acids whose entropy was below 1 (equivalent to 2 amino acids, having equal probability, so implying one amino acid present >50% of the time). Finally, we defined ZCWPW1 orthologues as those sequences matching at least 90% of those 31 perfectly conserved bases which were aligned, and aligning to at least 50% of these bases (this last condition allows for inclusion of incomplete sequencing or protein assembly). We note that while this 90% condition is arbitrary, by definition all annotated orthologues exceed this threshold, while thresholds below ∼70-80% are exceeded by orthologues of ZCWPW2 among other genes, offering some justification. Nonetheless, it is worth pointing out that our analysis does not rule out the existence of more poorly conserved ZCWPW1 copies in more distantly related species. This approach identified a final set of 167 genes, which we annotated as likely ZCWPW1 orthologues and used for the majority of results. For each ZCWPW1 orthologue, we also identified the taxonomic relationship of the closest species possessing such a PRDM9 copy (**Supplementary Table 1**). We also reciprocally annotated each PRDM9 copy previously identified (Baker et al., 2017) according to the closest species also possessing ZCWPW1 (**Supplementary Table 1**).

Our analyses revealed a relationship between ZCWPW1 predicted functional domains and PRDM9 histone modifications. Therefore, we identified additional potential conserved sites by identifying perfectly conserved bases among those species possessing *both* ZCWPW1 and PRDM9 orthologues, and where three key SET domain catalytic amino acids within PRDM9 are intact, meaning PRDM9 is predicted to have normal histone modification activity (Baker et al., 2017). This identified a slightly larger number of conserved amino acids (37). Only eight of these varied in any of the 167 species with a potential ZCWPW1 orthologue: 260, 404 and 411 (in two *Xenopus* frogs, with 411 also in white-headed capuchin), 19 and 22 (in three canines), 25 (in Ocelot gecko), 257 (in Anolis lizard and in Wombat), 325 (in elephantfish). While these changes might alter ZCWPW1 function, the true behaviour of ZCWPW1 is uncertain in these cases, though it is interesting that canids and frogs represent clades that all appear to have lost PRDM9, and possess multiple, clustered, amino acid changes.

### Domain Search

pDomThreader (Lobley et al., 2009), was used via the PSIPRED server (http://bioinf.cs.ucl.ac.uk/psipred/) on the following uniProt amino acid sequences (Q9H0M4|ZCPW1_HUMAN, Q6IR42|ZCPW1_MOUSE, E2RFJ2|E2RFJ2_CANLF, M3XJ39_LATCH, A0A3S5ZP38_BOVIN, M3WDY6_FELCA, and G3ULT5|G3ULT5_LOXAF) all of which identified a match to 1ub1A00 in the C terminal section after the PWWP domain, except in LOXAF in which this match was slightly below the p-value threshold of 0.001.

SCP1 prediction used NCBI conserved domain search server (https://www.ncbi.nlm.nih.gov/Structure/cdd/wrpsb.cgi?) with the uniProt amino acid sequences Q9H0M4|ZCPW1_HUMAN and Q6IR42|ZCPW1_MOUSE.

### Mice and genotyping

KO mice for *Zcwpw1* (C57BL/6N-Zcwpw1em1(IMPC)Tcp) were generated by the Toronto Centre for Phenogenomics (Canada). A 1485-bp deletion on chromosome 5 spanning exons 5 to 7 of *Zcwpw1* (chr5:137799545-13780101029) was engineered by CRISPR/Cas9 using guide RNAs 5’-GACTGCACTCACGGCCATCT-3’. The frameshift deletion introduces a stop codon in Exon 8, leading to a predicted unstable short truncated 492bp transcript. Mice were genotyped at the *Zcwpw1* locus using the following primers, and standard cycling conditions: KO allele-Forward, 5’-CACAGGCTCATGTATGTTTGTCTC-3’; KO allele-Reverse, 5’-CTGCTTCGTCCTCTTTCCTTATCTC-3’; WT allele-Forward, 5’-TGCCACCACACTTCATTTGT-3’; WT allele-Reverse CCTGTTTCCTTCCCAACTCA-3’. The deletion was verified by direct Sanger sequencing of the KO genotyping PCR product (Source Bioscience, UK), following purification with the QIAquick PCR Purification Kit (Qiagen). Sequence analysis was carried out using Chromas LITE (version 2.1.1).

KO mice for *Prdm9* were described previously (Hayashi et al., 2005; Mihola et al., 2019) and obtained from the RIKEN BioResource Research Center in Japan (strain name B6.129P2-Prdm9, strain number RBRC05145). Mice were genotyped at the *Prdm9* locus using the following primers, and standard cycling conditions: WT allele-Forward 5’-AGGAATCTTCCTTCCTTGCTGTCG-3’; WT allele-Reverse 5’-ATTTCCCTGTATCTTCTTCAGGACT-3’; KO allele-Reverse 5’-CGCCATTCAGGCTGCGCAACTGTT-3’.

All animal experiments received local ethical review approval from the University of Oxford Animal Welfare and Ethical Review Body (Clinical Medicine board) and were carried out in accordance with the UK Home Office Animals (Scientific Procedures) Act 1986.

### Fertility measurements

Fertility was assessed in mice ranging from 9 to 12 weeks of age, either by mating with WT littermates and recording the average litter size and frequency, or by measuring paired testes weight (normalized to lean body weight), sperm count (per paired epididymides) and chromosome synapsis rate (by immunostaining of pachytene spermatocytes) in males. Lean body weight was measured using the EchoMRI-100 Small Animal Body Composition Analyzer.

### Immunostaining of spermatocytes

Mouse testis chromosome spreads were prepared using surface spreading (Barchi et al., 2008; Peters et al., 1997) and immunostained as previously described (Davies et al., 2016) The following primary antibodies were used: custom ZCWPW1 rabbit antiserum (1:100), mouse anti-SYCP3 (Santa Cruz Biotechnology sc-74569, D-1) or biotinylated rabbit anti-SYCP3 (Novus NB300-232), rabbit anti-DMC1 (Santa Cruz Biotechnology sc-74569, D-1), rabbit anti-HORMAD2 (Santa Cruz Biotechnology sc-82192), mouse anti-RAD51 (Abcam ab88572), rabbit anti-RPA2 (Abcam ab10359), mouse (Millipore 05-636, clone JBW301) or chicken (Orbit orb195374, discontinued) anti-phospho γ-H2AX and Alexa Fluor 488-, 647- or 594-conjugated secondary antibodies against rabbit, mouse or chicken IgG (ThermoFisher Scientific), as well as avidin 647 (ThermoFisher Scientific) were used to detect the primary antibodies. Images were acquired using either a BX-51 upright wide-field microscope equipped with a JAI CVM4 B&W fluorescence CCD camera and operated by the Leica Cytovision Genus software, or a Leica DM6B microscope for epifluorescence, equipped with a DFC 9000Gt B&W fluorescence CCD camera, and operated via the Leica LASX software. Image analysis was carried out using Fiji (ImageJ-win64).

### FISH

Following immunostaining of spermatocytes, telomeres were labelled using the Telomere PNA FISH Kit/Cy3 (Agilent), following the manufacturer’s instructions, but without protease treatment. For the identification of chromosome 18, BAC RP24-144B10 (CHORI, BACPAC Resource Center) was labelled with the Abbot Molecular Nick Translation kit, and applied to the immunostained spermatocytes. The hybridisation and signal detection were carried out using standard techniques.

### Plasmids

Constructs encoding full-length human (h) or chimp (c) PRDM9 with a C-terminal V5 or N-terminal YFP tag for expression in mammalian cells (pLENTI CMV/TO Puro DEST backbone vector, Addgene plasmid # 17293; Campeau et al., 2009) were described previously (Altemose et al., 2017). Constructs encoding full-length human (h) and mouse (m) ZCWPW1 for expression in mammalian cells were purchased from Genescript (hZCWPW1 cDNA with a C-terminal HA tag in pCDNA3.1; clone ID OHu16813) and Origene (mZCWPW1 cDNA with a FLAG-Myc dual tag in pCMV6-Entry; clone ID MR209594), respectively. For expression in *E. Coli*, full-length mZCWPW1 cDNA was subcloned into the pET22b(+) vector (Novagen) with a C-terminal poly-His tag.

### ZCWPW1 antibody production

pET22b-mZCWPW1-His construct was transformed into BL21 (DE3) *E.Coli* cells (Invitrogen). Bacterial cultures were grown to a density with O.D_600_∼0.7, and expression of recombinant His-tagged ZCWPW1 protein was induced overnight at 20°C by addition of IPTG to 0.5mM. The protein was purified using Talon metal affinity resin, according to the manufacturer’s instructions (Takara). Further purification was carried out by size exclusion chromatography (Superdex HiLoad 200 16/60, GE Life Sciences). Western Blot validation was carried out as described previously (Altemose et al., 2017), using mouse anti-human ZCWPW1 (SIGMA SAB1409478) and mouse anti-polyhistidine (SIGMA H1029). The purified protein was used to immunize 2 rabbits (Eurogentec, Belgium), and the resulting immune antisera were tested against the recombinant antigen by ELISA (Eurogentec), and the endogenous protein in mouse testes from WT and KO mice (data not shown), alongside the preimmune sera.

### Cell line, transfection and immunofluorescence staining

Human embryonic kidney (HEK) 293T cells were cultured and transfected using Fugene HD as previously described (Altemose et al., 2017). High and comparable expression of the target proteins was verified by immunofluorescence staining of duplicate transfected cultures before proceeding to ChIP. PRDM9 expression was visualized either directly from live cells through YFP fluorescence, or by immunofluorescence staining of fixed cells using rabbit anti-V5 (Abcam ab9116) as described previously (Altemose et al., 2017). ZCWPW1-HA was detected by immunostaining as above, using rabbit anti-HA (Abcam ab9110, 1:100). The fraction of cells co-expressing ZCWPW1-HA and PRDM9-V5 was determined by co-staining with both antibodies, as above (**Supplementary Figure 11**).

### ChIP-seq

DMC1 ChIP-seq data from WT B6 mouse testis were generated previously (Brick et al., 2012). H3K4me3 ChIP-seq data was generated previously (Davies et al., 2016).

#### ChIP

ChIP against ZCWPW1-HA, (h/cPRDM9-V5) was carried out from transfected HEK293T cells as previously described (Davies et al., 2016); (Altemose et al., 2017) were crosslinked for 10min in 1% formaldehyde, the reaction was quenched for 5min by the addition of glycine to a final concentration of 125mM, and the cells were washed twice in cold PBS. The cell pellet was resuspended in cold sonication buffer (50mM Tris-HCl pH8, 10mM EDTA, 1% SDS) supplemented with protease inhibitors (Complete, Roche), and chromatin was sheared to an average size of 200-500bp by sonication for 35 cycles (30s ON/30s OFF) using a Bioruptor Twin (Diagenode). After centrifugation for 10min at 20,000g, 4°C, the sonicate was pre-cleared for 2h at 4°C with 65μl of Dynabeads™ M-280 sheep anti-rabbit IgG (Life Technologies) and a 1% input chromatin sample was set aside. The rest of the sample was diluted 10 fold in ChIP buffer (16.7mM Tris pH8 1.2mM EDTA, 167mM NaCl, 1.1% Triton X-100) supplemented with protease inhibitors, and incubated overnight at 4°C with 5μl of either Abcam rabbit antibody: anti-HA (ab9110), anti-H3K4me3 (ab8580), anti-H3K36me3 (ab9050). Immunocomplexes were washed once with each of low salt buffer (20 mM Tris pH8, 150 mM NaCl, 1% Triton X-100, 0.1% SDS, 2mM EDTA), high salt buffer (20 mM Tris pH8, 500 mM NaCl, 1% Triton X-100, 0.1% SDS, 2mM EDTA), LiCl buffer (10mM Tris pH8, 0.25M LiCl, 1% NP-40, 1% sodium deoxycholate, 1mM EDTA) and TE buffer (10mM Tris pH8, 1mM EDTA) and eluted from the beads in 100 mM NaHCO_3_, 1% SDS for 30 min at 65°C with shaking. Both input and ChIP samples were reverse crosslinked overnight at 65°C in the presence of 200mM NaCl, and proteins were digested for 90 min at 45°C by addition of proteinase K (0.3ug/ul final concentration). DNA was purified using the MinElute reaction cleanup kit (QIAGEN), and quantified using the dsDNA HS assay kit and a Qubit® 2.0 Fluorometer (Invitrogen).

ChIP against DMC1 was performed from *Zcwpw1^−/−^* testes using the published method by (Khil et al., 2012) with some modifications listed here. Chromatin shearing was carried out in 20mM Tris-HCl pH8, 2mM EDTA, 0.1% SDS using a Bioruptor Pico sonicator (Diagenode) for 4 cycles of 15s ON/45s OFF. ChIP was performed in 10mM Tris-HCl pH8, 1mM EDTA, 0.1% Sodium Deoxycholate, 1% Triton X-100, 500mM NaCl using 5μg of mouse anti-DMC1 2H12/4 (Novus NB100-2617) pre-bound to 50μl of Dynabeads™ M-280 sheep anti-mouse IgG (Life Technologies).

#### Sequencing

ZCWPW1 ChIP and input libraries from transfected cells were prepared by the Oxford Genomics Centre at the Wellcome Centre for Human Genetics (Oxford, UK) using the Nextera DNA Library Prep Kit and established Illumina protocols, and sequenced on a HiSeq 4000 platform (75bp paired end reads, 48 million reads/sample). DMC1 ChIP libraries from *Zcwpw1^−/−^* testes were prepared and sequenced as described previously (Davies et al., 2016) on an Illumina HiSeq2500 platform (Rapid Run, 51bp paired end reads, 110 million reads/sample).

### Read Mapping

For the HEK293 experiments, reads were mapped to either hg38 (NCBI’s GCA_000001405.15_GRCh38_no_alt_plus_hs38d1_analysis_set.fna.gz) using bwa mem (version 0.7.17-r1188) (Li, 2013). Duplicates were removed using picard’s markDuplicates (version 2.20.4-SNAPSHOT). Unmapped, mate unmapped, non primary alignment, failing platform, and low MAPQ reads were removed using samtools with parameters “-q 30 -F 3852 -f 2 -u” (version 1.9 (using htslib 1.9)). Other unmapped reads and secondary alignments were removed using samtools fixmate. Fragment position bed files were created using bedtools bamtobed (v2.28.0).

For the DMC1 mouse experimental data, we processed the data following the algorithm provided by (Khil et al., 2012) to map the reads to the mouse mm10 reference genome (Lunter and Goodson, 2011), and obtain type I reads.

### Peak Calling

We called DMC1 peaks, as described previously (Davies et al., 2016). For the HEK293 experiments, peaks for ZCWPW1, PRDM9, H3K4me3, and H3k36me3 were called using a peak caller previously described (Davies et al., 2016) and available at https://github.com/MyersGroup/PeakCaller. Single base peaks were called with parameters pthresh 10^−6^ and peakminsep 250 on 22 autosomes. For each experiment, we used available IP replicates, and sequenced input DNA to estimate background, in calling peaks. Most of our analyses refer to these peaks, except that Figures 5A and B plot the *change* in ZCWPW1 occupancy following PRDM9 transfection, for either the chimp or human PRDM9 alleles. To estimate ZCWPW1 levels with vs. without PRDM9, we called ZCWPW1 peaks in HEK293T cells also transfected with PRDM9 as before, but now replacing the “input” lane with the IP data for ZCWPW1 in cells not transfected with PRDM9. **Supplementary Figure 13** also uses this measure, of ZCWPW1 recruitment attributable to PRDM9 binding, to predict DMC1 levels, while **Supplementary Figure 21** (right panel) checks that changes in ZCWPW1 occupancy occur largely at PRDM9-bound sites. We analysed previously published H3K4me3 and SPO11 read data (Davies et al., 2016; Lange et al., 2016).

### Enrichment Profiles for HEK293T experiments

Peaks were filtered if: center within 2.5kb of PRDM9 independent H3K4me3 (promoters), input coverage <=5, in top 5 by likelihood, greater than 99.9 percentile input coverage. For human PRDM9 peaks the PRDM9 motif position (Altemose et al., 2017) was inferred using the getmotifs function in MotifFinder within the 300bp region around the peak (with parameters alpha=0.2, maxits=10, seed=42, stranded_prior=T), and peaks were recentered and stranded at these locations. Peaks within 4kb of one another are removed to avoid double counting. Mean coverage was calculated with bwtool 1.0 using the aggregate command with parameters “-fill=0 -firstbase” at a width of +/−2kb. Each profile was normalised by the total coverage of all of the fragments as well as by the input profile. Random profiles were created using bedtools random with 10,000 locations, with seed 72346, and were also filtered such that non were within 4kb of one another.

### DSB Profiles

Mapped type 1 reads were filtered by removing non-canonical chromosomes (with underscores in names), sex chromosomes, and mitochondrial reads. Bedgraphs were converted to bigWig format using UCSC bedGraphToBigWig v4. Profiles were created using bwtool 1.0 using the aggregate command with parameters 5000:2000 and -fill=0. B6 wild type hotspot locations were filtered to remove non-autosomal chromosomes, retain only B6 allele hotspots, and remove hotspots with PRDM9-independent H3K4me3. *Prdm9*^−/−^ hotspots were filtered to remove X chromosome. Motif centered coordinates were used for Wild type hotspots. To normalise background signal, the mean signal between −5000 and −3000 was subtracted from each strand-mouse combination. Mean coverage was normalised by the sum of coverage over each strand-mouse combination across both WT and KO locations.

### Mapping of Alu CpGs

Alu locations were downloaded from UCSC tables and filtered for Alu repeats with a width between 250 and 350bp. DNA sequence was extracted from the genome (Ensembl 95 h38 primary assembly) at these locations using the bedtools getfasta command and CpG dinucleotides were counted using ‘stringr::str_locate_all’ command in R.

### DMC1 Prediction using ZCWPW1 and PRDM9 binding strength

PRDM9-dependent ZCWPW1 was force-called at the human PRDM9 peaks. DMC1 sites are from (Pratto et al., 2014) (GSE59836) and were subsetted to ‘AorB’ autosomal peaks with a width less than 3kb. These peaks were then trimmed to a maximum of 800bp before overlapping with PRDM9 peaks to create a binary target variable. Regions with input coverage <=5 in either predictor were removed in addition to outliers with input > 200 and or Zcwpw1 enrichment >10. Chromosomes 1,3, and 5 were used as the test data with the remaining autosomes being the training data.

### Heatmaps

Regions were extracted from bigWig using bwtool matrix -fill=0 -decimals=1 -tiled-averages=5, and width parameters 2000. Coverage was normalised by total coverage (scaled by 10^10^). A pseudocount of 1 was added to both input and sample, and the sample was then normalised by the input for each region. Values outside the quantile range 0.01-0.99 were thresholded. The profile plots are created by taking the ratio of the mean coverage of the sample and input separately. Ordering of the regions was determined by the mean coverage of a 200bp window centered on the peak center.

## Acknowledgments

We thank the High-Throughput Genomics Group at the Wellcome Centre for Human Genetics (funded by Wellcome Trust grant 203141/Z/16/Z) for the generation of sequencing data. We also thank Dr. Benjamin Bishop and Prof. Christian Siebold in the Division of Structural Biology (Wellcome Centre for Human Genetics, Oxford) for their help with the purification of recombinant mouse ZCWPW1 (size exclusion) towards antibody production. Funding: This work was supported by a Wellcome Trust Investigator Award to S.R.M. (098387/Z/12/Z & 212284/Z/18/Z) and Wellcome Trust PhD Studentship to D.W. (109109/Z/15/Z). Graphical abstract was created with BioRender.com.

## Author contributions

S.R.M. designed the study; E.B carried out cloning, cell culture experiments, bred the mice and collected fertility measures; D.M. performed immunostaining of spermatocytes and FISH for protein expression and synapsis analysis; E.B performed ZCWPW1 ChIP-seq in human cells and G.Z. performed DMC1 ChIP-seq in mouse testis; E.B, D.W, D.M, A.G.H (DMC1 ChIP-seq) and S.R.M analysed the data; E.B, D.W and S.R.M prepared the manuscript.

## Materials & Correspondence

Code is available at www.github.com/MyersGroup/Zcwpw1. Other requests should be addressed to S.R.M.

