## Supplementary_Figures_&_Tables_2-4 for "ZCWPW1 is recruited to recombination hotspots by PRDM9, and is essential for meiotic double strand break repair"

### Supplement

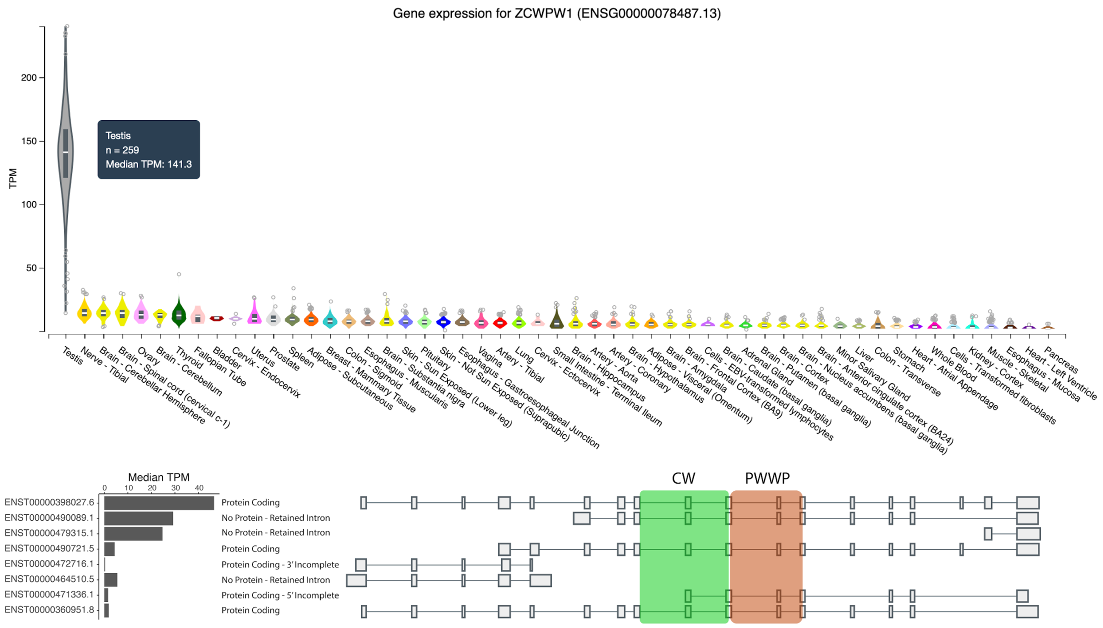


**Supplementary Figure 1**. *ZCWPW1* is specifically expressed in testis in humans. Data Source: GTEx Analysis Release V7 (dbGaP Accession phs000424.v7.p2). Top: Total expression by human tissue type. Bottom: Isoforms of *ZCWPW1* expressed in human testis.

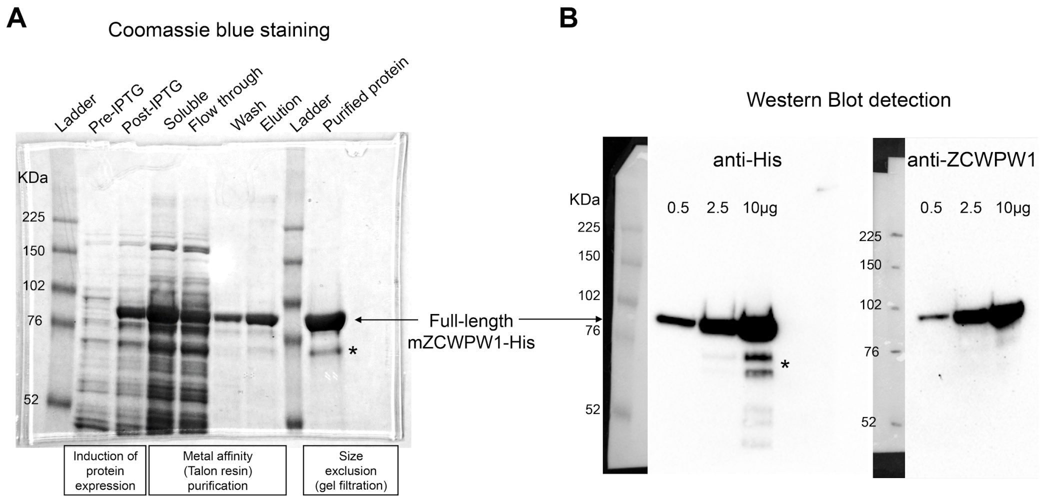


**Supplementary Figure 2**. Purification of recombinant full-length mouse ZCWPW1 protein. (A) SDS-PAGE analysis and Coomassie blue staining of bacterial lysates before (pre-IPTG) and after (post-IPTG) induction of protein expression with IPTG, the soluble protein fraction (after cell sonication) used for purification, the flow through (incompletely depleted from the target protein) after incubating the soluble fraction with Talon resin beads to bind His-tagged mZCWPW1, the wash containing 5mM imidazole, the protein eluate from the beads using 300mM imidazole, and the purified recombinant protein after further purification from low MW contaminants by size exclusion. (B) Western blot detection of purified His-tagged mZCWPW1 using an anti-His and a mouse polyclonal antibody raised against the human protein (previously tested positively against mouse ZCWPW1 overexpressed in HEK293T cells). *indicates degradation fragments (likely C-terminal).

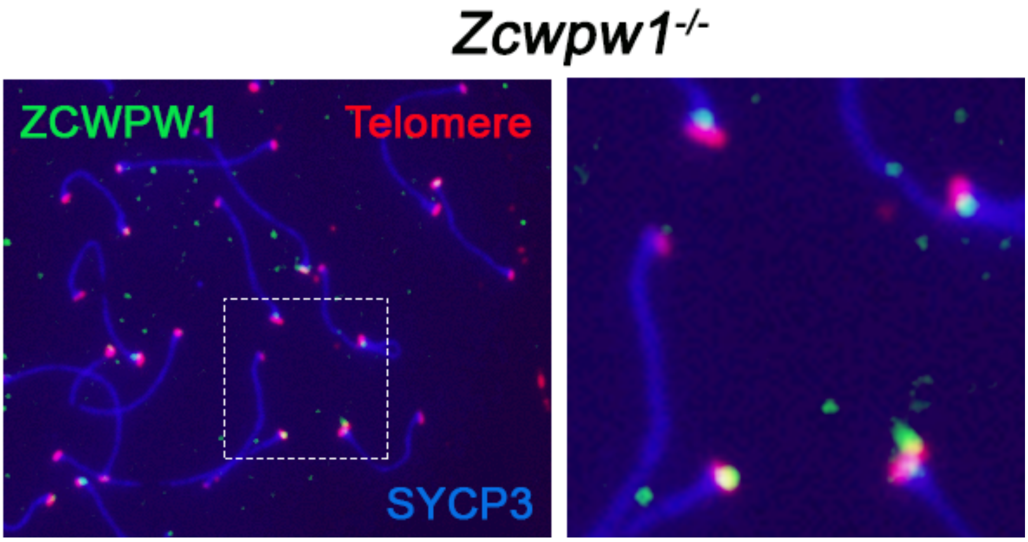


**Supplementary Figure 3**: ZCWPW1 localises at subtelomeric regions of chromosomes in pseudo-pachytene cells. Testis chromosome spreads from *Zcwpw1*$^{-/-}$ were immunostained for SYCP3 and ZCWPW1, and hybridised by FISH with a telomeric probe. Note that the ZCWPW1 foci do not always exactly co-localize with the telomeric signal, and generally lie more internally on the synaptonemal complex axis. Right panel: 3x zoom from left image showing the area in the dashed white rectangle.

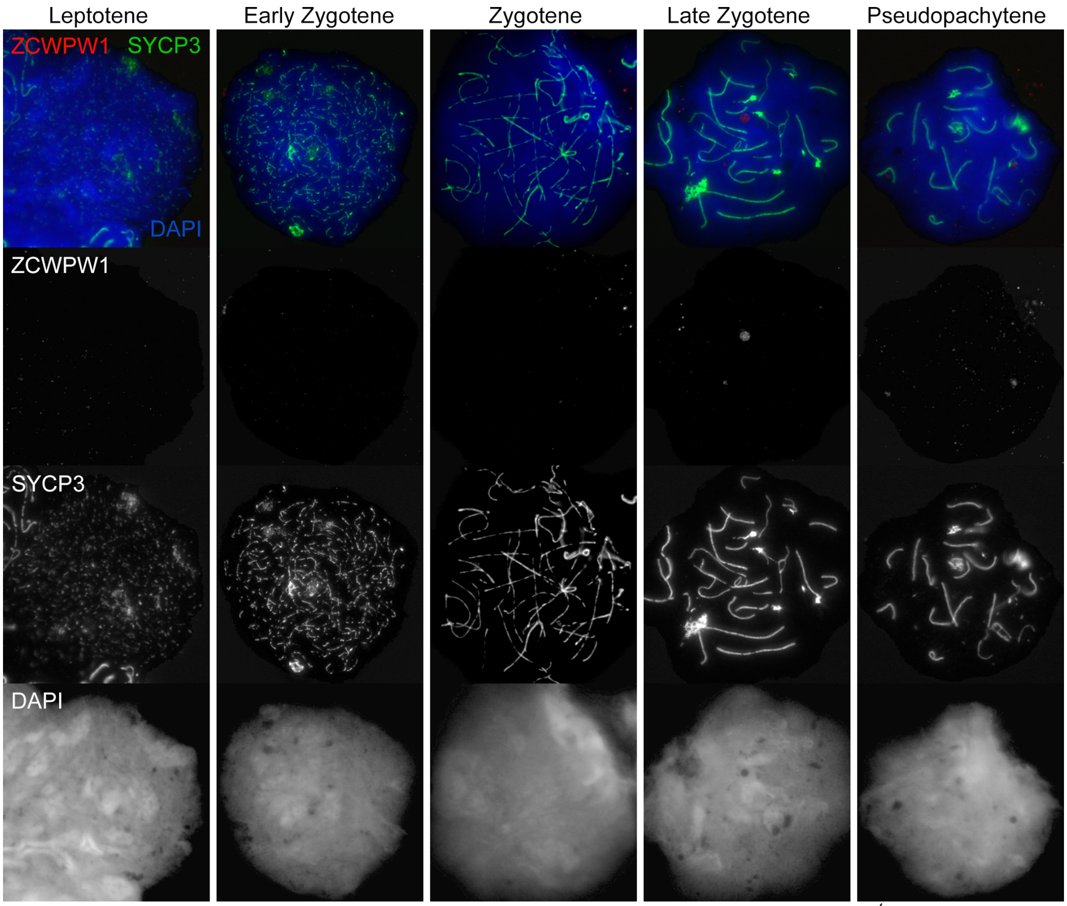


**Supplementary Figure 4.** Loss of ZCWPW1 expression in *Zcwpw1*$^{-/-}$ mouse testis. Testis chromosome spreads from *Zcwpw1*$^{+/+}$ and *Zcwpw1*$^{-/-}$ mice were immunostained with antibodies against the synaptonemal complex protein SYCP3, and ZCWPW1, and counterstained with DAPI to visualise nuclei. Developmental stages are indicated at the top. The top row of panels shows merged signals, and the bottom raw individual signals.

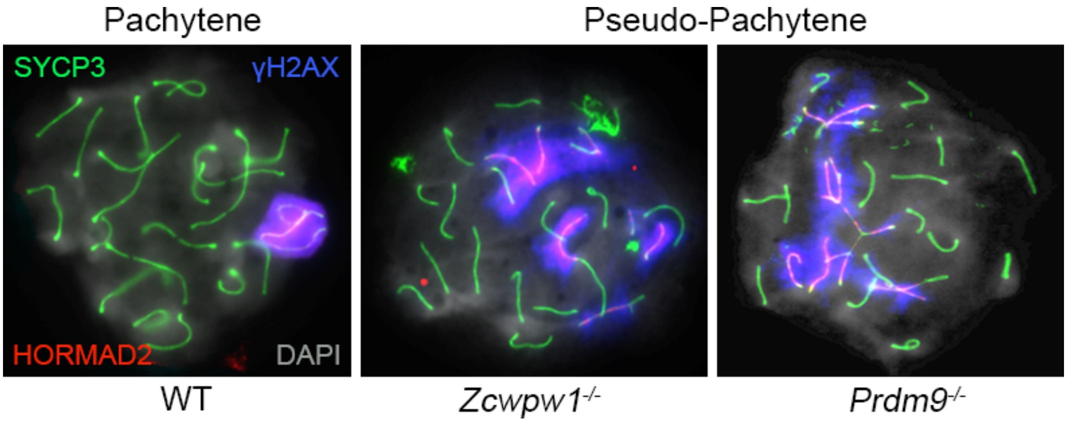


**Supplementary Figure 5.** Asynapsis and lack of XY body formation in *Zcwpw1*$^{-/-}$ mouse testis. Testis chromosome spreads from WT, *Zcwpw1*$^{-/-}$ and *Prdm9*$^{-/-}$ mice were immunostained with antibodies against SYCP3, γ-H2AX (phosphorylated form) and HORMAD2, and counterstained with DAPI. WT Pachytene cells show full synapsis of all autosomes and an XY body strongly labelled with γ-H2AX. In contrast, the XY body is absent in *Zcwpw1*$^{-/-}$ and *Prdm9*$^{-/-}$ pseudo-Pachytene cells, with no clear XY body formation. In the *Prdm9*$^{-/-}$ mutant, mispairing of homologs is evident by the formation of branched structures referred to as “tangled” chromosomes in the text and **Supplementary Table 4**.

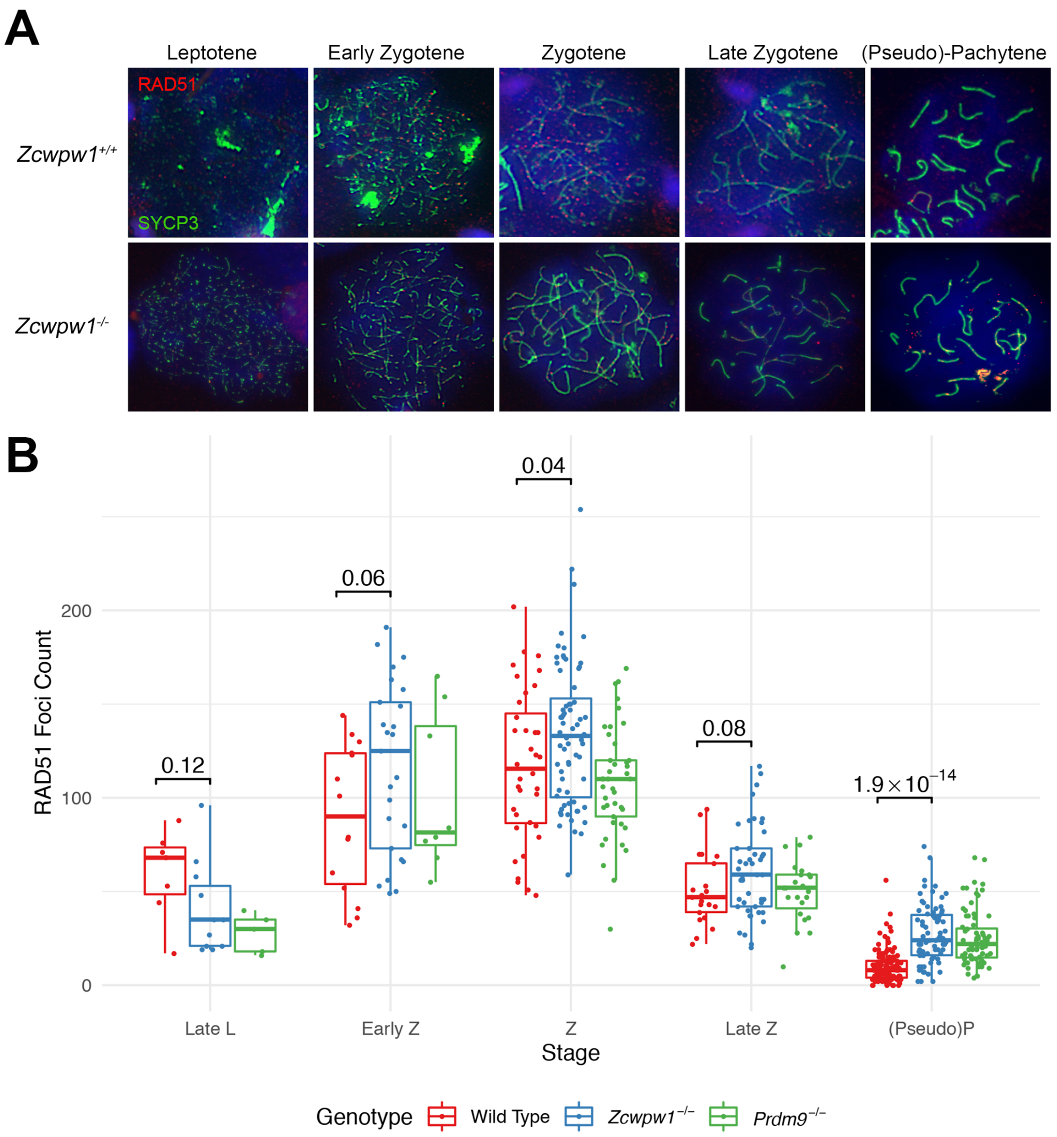


**Supplementary Figure 6.** (**A**) RAD51 staining in *Zcwpw1*$^{-/-}$ mouse testis. Testis chromosome spreads from *Zcwpw1*$^{+/+}$ and *Zcwpw1*$^{-/-}$ mice were immunostained with antibodies against the synaptonemal complex protein SYCP3 and the recombinase RAD51, and counterstained with DAPI to visualise nuclei. Developmental stages are indicated at the top. (**B**) The number of RAD51 foci in cells from the various stages of prophase I were counted. p-values are from Welch's two sided, two sample t-test. L: Leptotene, Z: Zygotene, P: Pachytene. n=2 mice per genotype (*Zcwpw1*$^{-/-}$ and WT), n=1 for *Prdm9*$^{-/-}$.

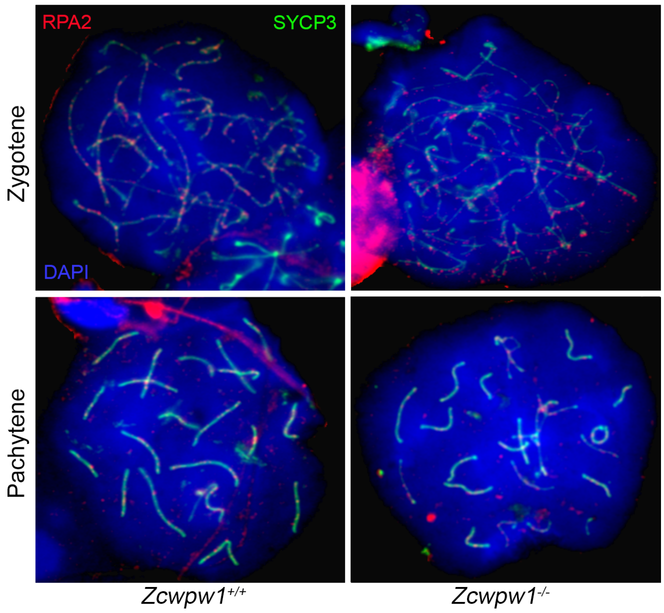


**Supplementary Figure 7.** RPA2 staining in *Zcwpw1*$^{-/-}$ mouse testis. Testis chromosome spreads from *Zcwpw1*$^{+/+}$ and *Zcwpw1*$^{-/-}$ mice were immunostained with antibodies against SYCP3 and RPA2, and counterstained with DAPI to visualise nuclei. Developmental stages are indicated on the side.

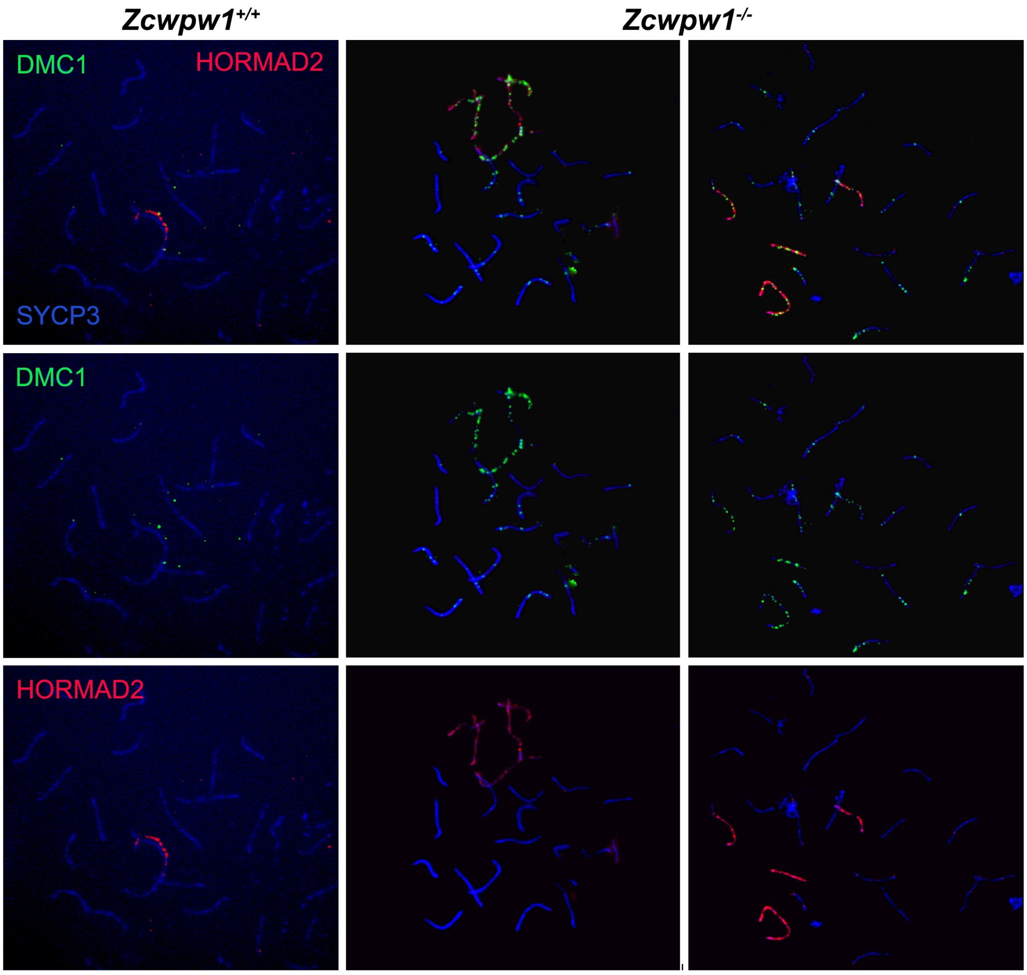


**Supplementary Figure 8.** DSB repair is delayed with accumulation of DMC1 on asynapsed chromosomes in the *Zcwpw1*$^{-/-}$ mouse. Testis chromosome spreads from *Zcwpw1*$^{+/+}$ *and Zcwpw1*$^{-/-}$ mice were immunostained for DMC1, HORMAD2, and SYCP3. Representative images of 2 mutant pseudopachytene cells show accumulation of DMC1 foci on HORMAD2-positive asynapsed chromosomes. In contrast wild-type pachytene cells only show residual DMC1 foci on partially synapsed XY sex chromosomes. Note that in mutant cells, even synapsed chromosomes abnormally retain some level of DMC1.

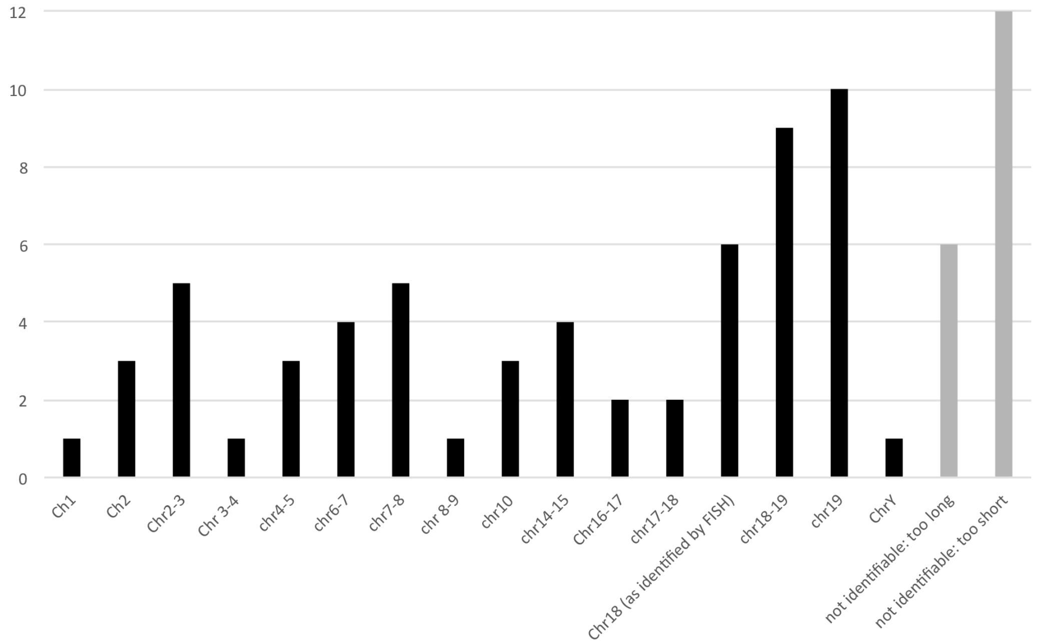


**Supplementary Figure 9**. Frequency of asynapsis per chromosome in *Zcwpw1*$^{-/-}$ males. Testis chromosome spreads were immunostained for SYCP3, HORMAD2 and γ-H2AX to identify asynapsed chromosomes in pseudopachytene cells, and hybridized with a probe specific to chromosome 18 (chr18). The identity of the other chromosomes in the same cell was determined using the correlation between the length of the SYCP3 signal (in pixels) for chr18 and the known size of this chromosome (in Mb) as a pixel to Mb ratio reference. The frequency of each chromosome, or group of chromosomes of similar length which the analysis cannot tell apart, was plotted. Number of informative cells analysed n=16 (only fully asynapsed chromosomes were considered in the analysis).

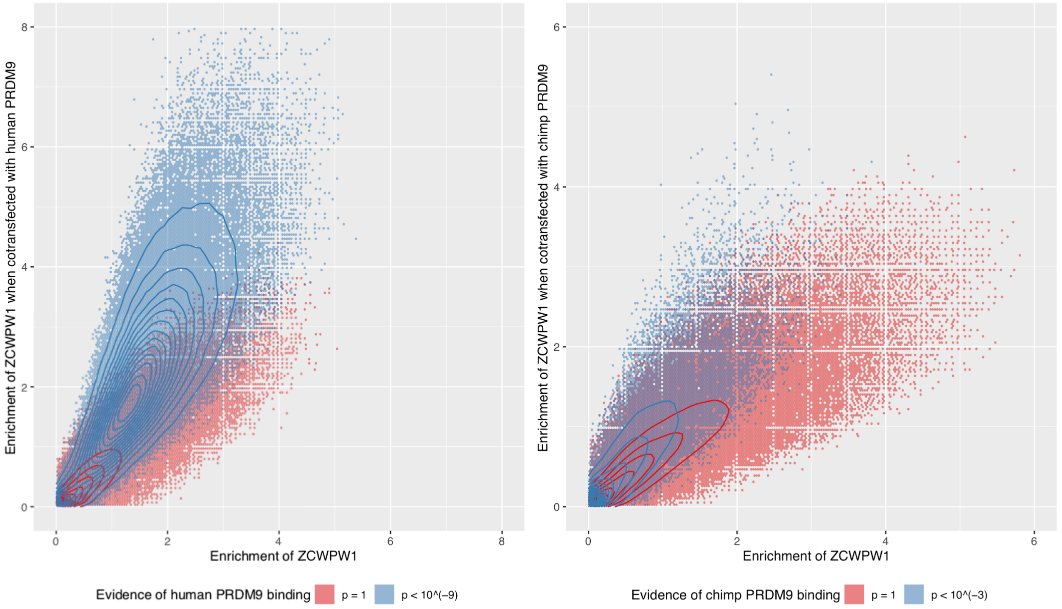


**Supplementary Figure 10**. Enrichment of ZCWPW1 when co-transfected with PRDM9 is dependent on the ability of ZCWPW1 to bind in the absence of PRDM9. Enrichment was force called in 100bp windows across the genome. Data is conditioned on having input coverage of greater than 5 and enrichment>0.01 for both axes. Hexagons are coloured if at least 3 data points are present. Solid lines show density contours estimated by MASS::kde2d() in R.

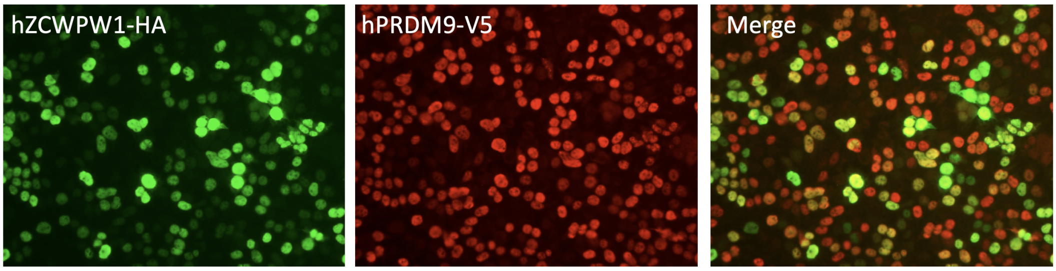


**Supplementary Figure 11**. Co-transfection of ZCWPW1 and PRDM9 in HEK293T cells. Immunofluorescence staining against the protein tags shows high expression levels of each protein and a reasonable proportion of co-expressing cells with merged overlapping signals (ranging from light green to yellow and light red depending on the expression ratio of the two proteins).

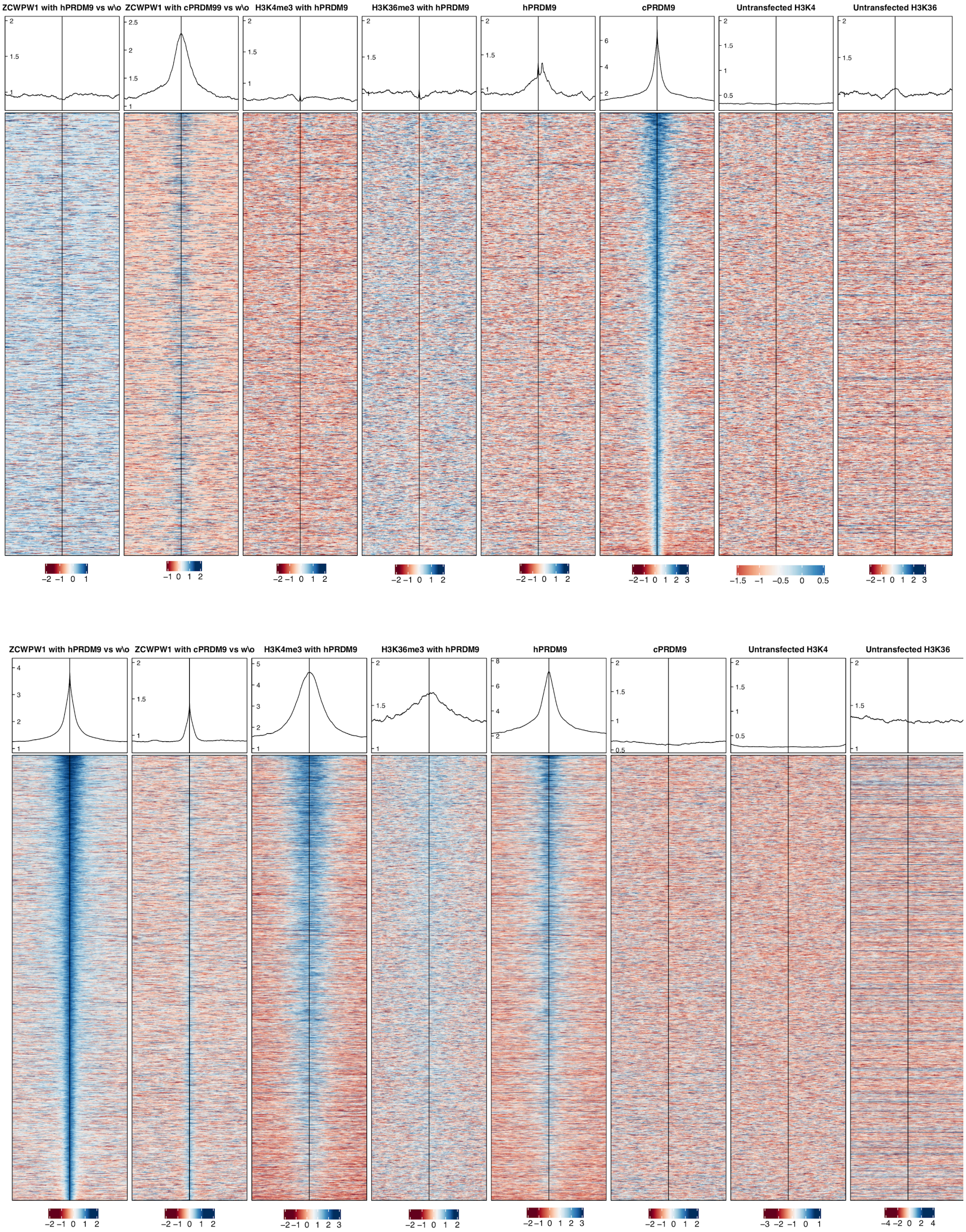


**Supplementary Figure 12. (A)** Profiles and heatmaps of reads at locations of chimp PRDM9. Heatmaps show log fold change of sample (as indicated in the title of each column, methods) vs input, for the top ¼ of peaks of chimp (c) PRDM9, for various samples, ordered by chimp PRDM9. Here showing that ZCWPW1 is found at sites of chimp PRDM9 peaks, when co-transfected with chimp PRDM9, but not at human (h) PRDM9 peaks. **(B)** Profiles and heatmaps of reads at locations of ZCWPW1 co-transfected with human PRDM9. Heatmaps show log fold change of sample (as indicated in the title of each column, Methods) vs input, for the top ¼ of peaks of ZCWPW1 when co-transfected with PRDM9, for various samples, ordered by first column. Here showing that H3K4me3, H3K36me3 and hPRDM9 are found at ZCWPW1 peaks when co-transfected with PRDM9.

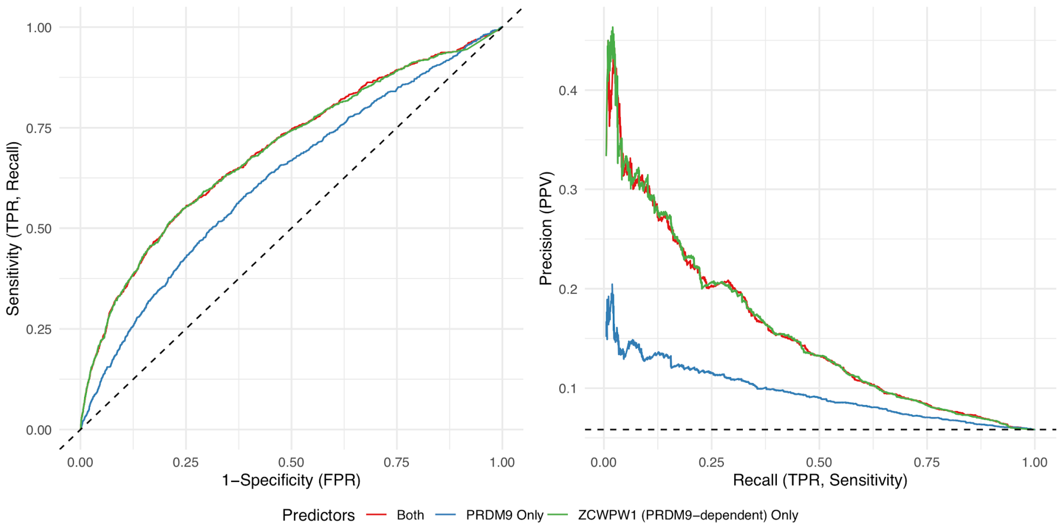


**Supplementary Figure 13:** ZCWPW1 enrichment (with PRDM9 vs without) provides a better predictor of DMC1 sites than PRDM9 itself using a logistic regression (Methods). **(A)** Receiver Operating Characteristic curve **(B)** Precision Recall Curve. ZCWPW1 enrichment (with PRDM9 vs without) refers to enrichment of ZCWPW1 cotransfected with PRDM9 relative to (using as input) ZCWPW1 transfected alone.

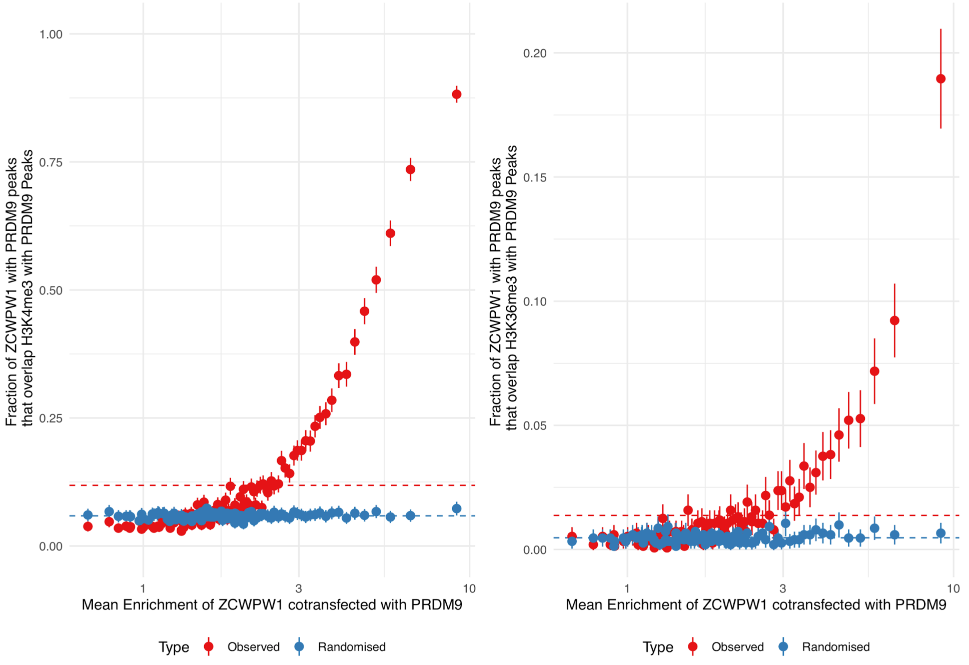


**Supplementary Figure 14.** Fraction of ZCWP1 peaks (co-transfected with PRDM9 with input coverage of at least 5) that overlap either H3K4me3 or H4K36me3 for different bins of ZCWPW1 enrichment (100 equal sample size bins). Error bars show ±2 s.e. of the proportion. ‘Randomised’ shows expected proportions when x-axis regions are randomly shifted within a range of 1e8 bases (up to a maximum of the length of the chromosome).

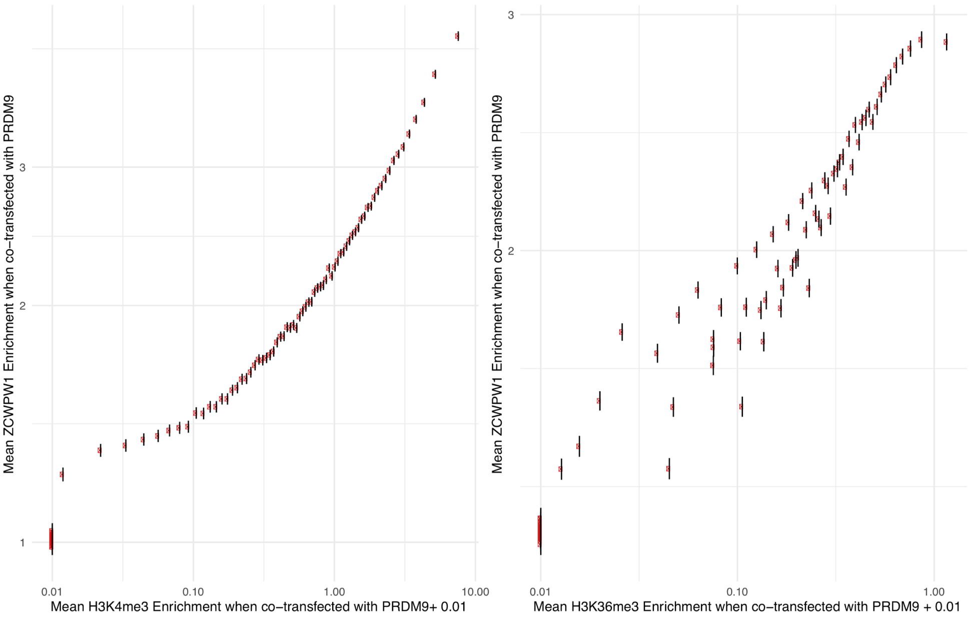


**Supplementary Figure 15.** Enrichment from 100bp non-overlapping windows is binned into 100 equal sample size bins and mean enrichment of ZCWPW1 cotransfected with PRDM9 is plotted. Error bars show ±2 s.e. of the mean. Windows with evidence of PRDM9-independent H3K4me3 have been removed from the H3K4me3 plot. Additionally, x-axis regions were removed if input reads were <15 and y-axis regions if <5.

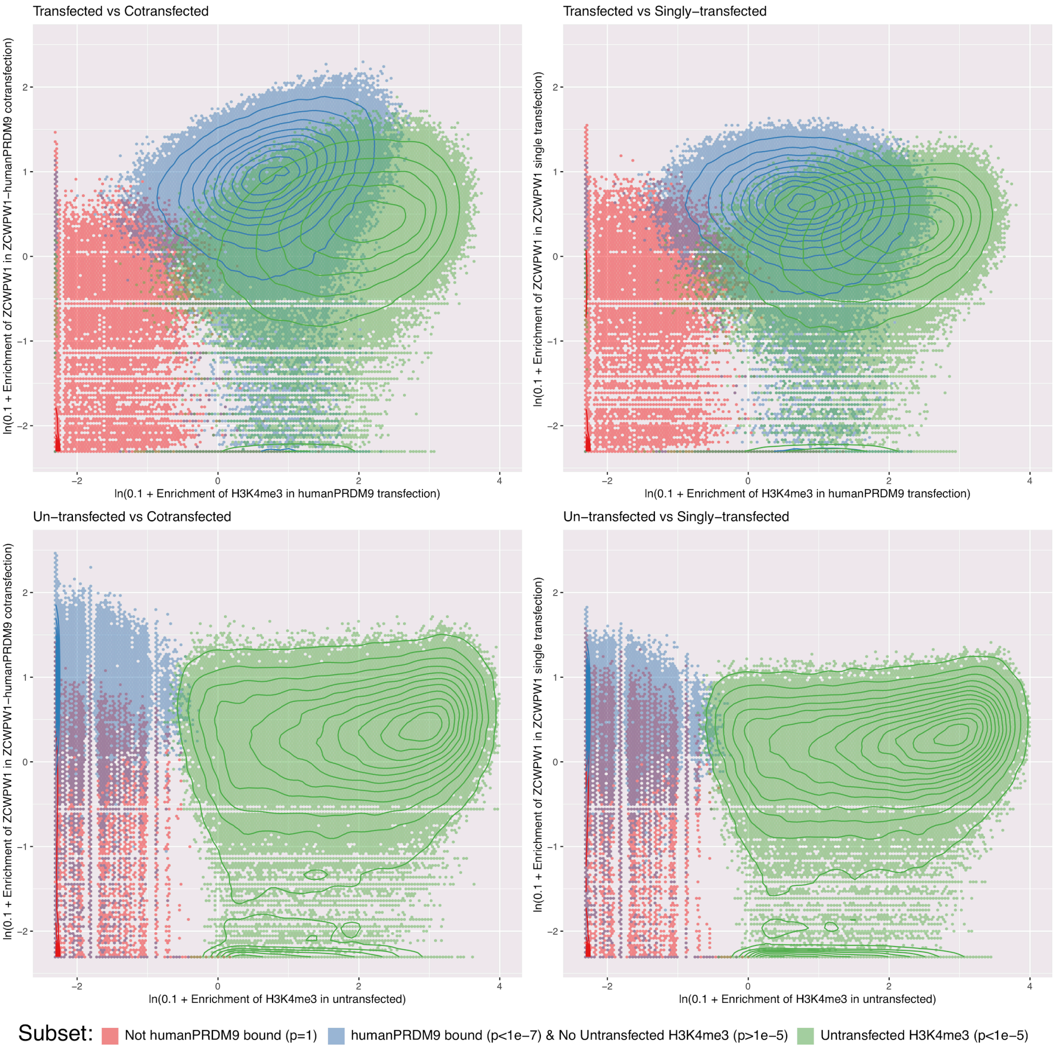


**Supplementary Figure 16. Dependence of ZCWPW1 enrichment on H3K4me3.** Subplot titles describe human PRDM9 transfection status vs ZCWPW1 cotransfection (with human PRDM9) status. Enrichment was force called in 100bp windows across the genome. Data for each colour group was downsampled to have the same number of data points per group. Hexagons are coloured if at least 5 data points fall within that bin. Solid lines show density contours estimated by MASS::kde2d() in R.

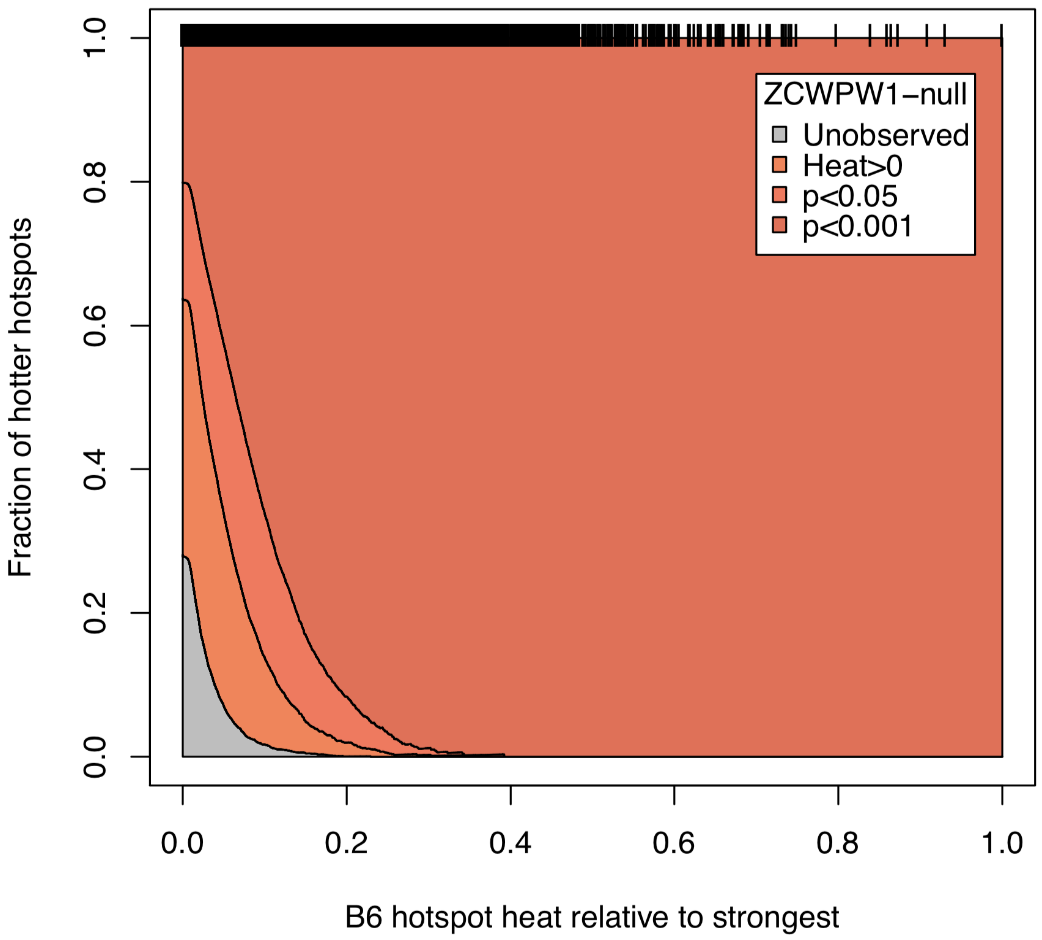


**Supplementary Figure 17.** Fraction of B6 hotspot locations seen in *Zcwpw1*$^{-/-}$ DMC1 ChIP-seq at different p-values. Heat refers to DMC1 signal. Black rung bars across the top each represent a single peak. We observe almost all of the peaks seen in B6 wild type, especially for the hottest hotspots.

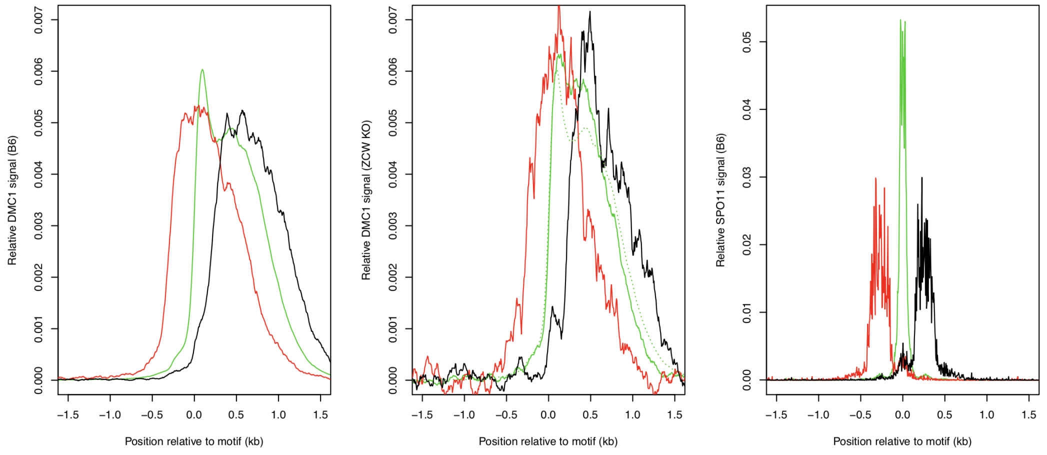


**Supplementary Figure 18. DSBs in *Zcwpw1*$^{-/-}$ are positioned at WT locations within hotspots.** DMC1 signal (left and center) is stratified by SPO11 (right). Stratification is into active hotspots (top 30%) with >90% of the SPO11 signal in the central 300bp (green), and <50% central, and >90% upstream of the PRDM9 binding motif (red) or <10% upstream (black).

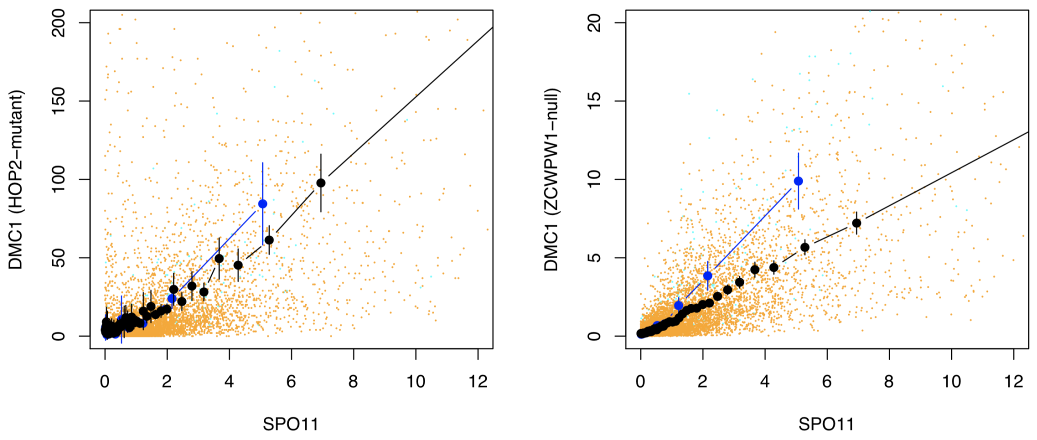


**Supplementary Figure 19.** *Hop2*$^{-/-}$ and *Zcwpw1*$^{-/-}$ mouse knockout mutants show the same linear relationship of DMC1 ChIP-seq vs SPO11 (WT). DMC1 data from *Hop2*$^{-/-}$ mice is from GSM851661 (Khil et al., 2012).

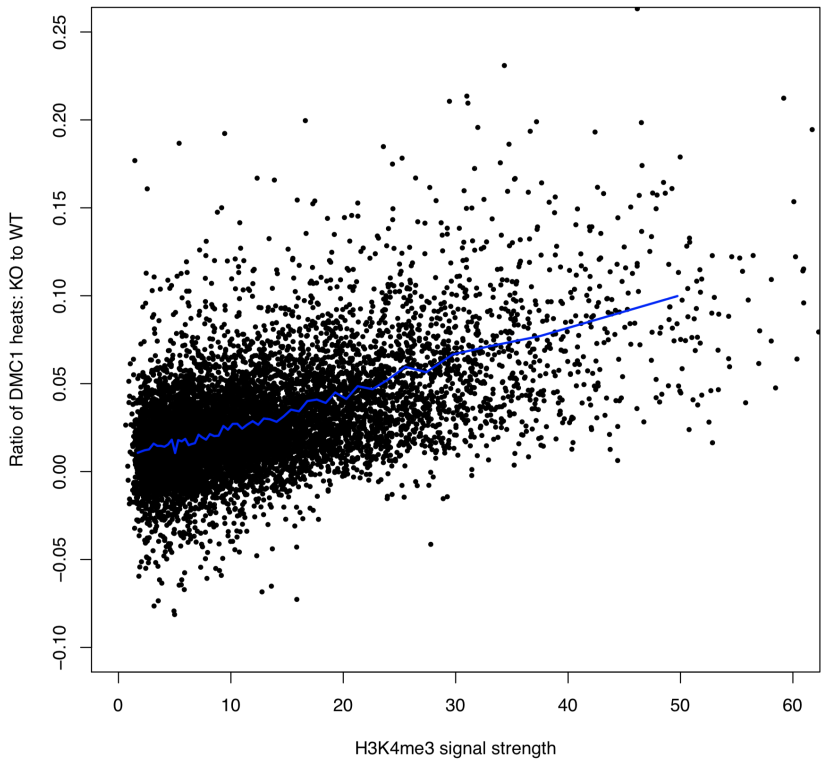


**Supplementary Figure 20.** Regression of DMC1 ratio on H3K4me3. For each B6 mouse hotspot (Methods), we calculated the ratio of DMC1 fragment coverage (summed over the central 3.5kb region spanning DMC1 hotspot enrichment in *Zcwpw1*$^{-/-}$ and WT mice, after subtracting mean background over the ±2.5-5kb distal regions, and removing any hotspot with mean SPO11 coverage over the surrounding ±5kb outside the hotspot >1, to exclude nearby hotspots). We excluded weak hotspots whose estimated SPO11 heat (within 2.5kb) or DMC1 WT heats were in the bottom 33% (because accurate ratio estimation is not possible for these hotspots). Circles: the force-called H3K4me3 signal strength, vs. the ratio, for each of 9,318 resulting hotspots. Blue line: mean H3K4me3 signal versus the ratio of mean DMC1 fragment coverages, after binning by H3K4me3 signal into 50 equal sample size groups of hotspot, within each bin. The points and blue line show a strong and essentially linear relationship.

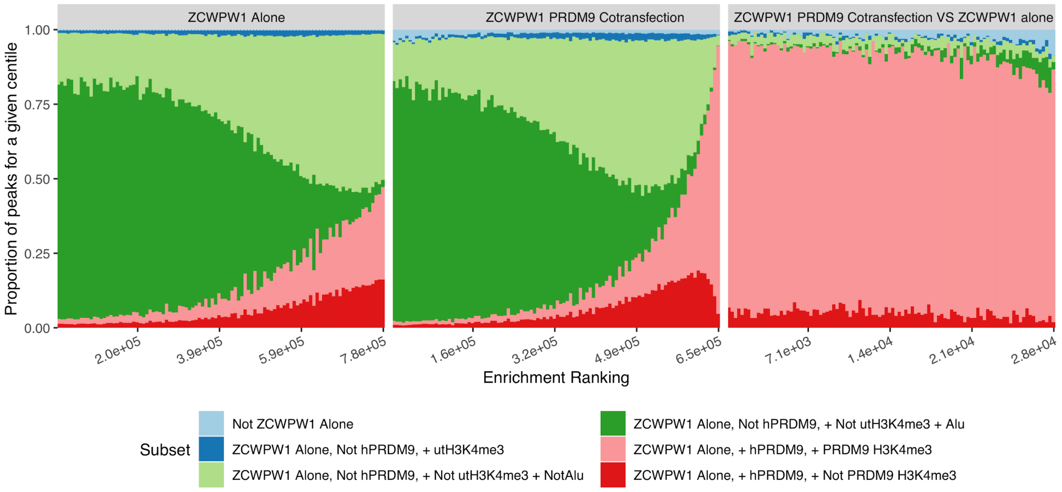


**Supplementary Figure 21.** Proportion of ZCWPW1 peaks overlapping various other marks, ordered by enrichment of ZCWPW1 binding over input. hPRDM9: human PRDM9; ut: untransfected. Subset categories are labelled according to the following example: "ZCWPW1 alone, Not hPRDM9, + Not utH3K4me3 + Not Alu" means ZCWPW1 binding peaks in cells transfected with ZCWPW1 alone that do not correspond to human PRDM9 binding peaks, or H3K4me3 peaks (ChIP-seq data in untransfected cells), or Alu repeats.

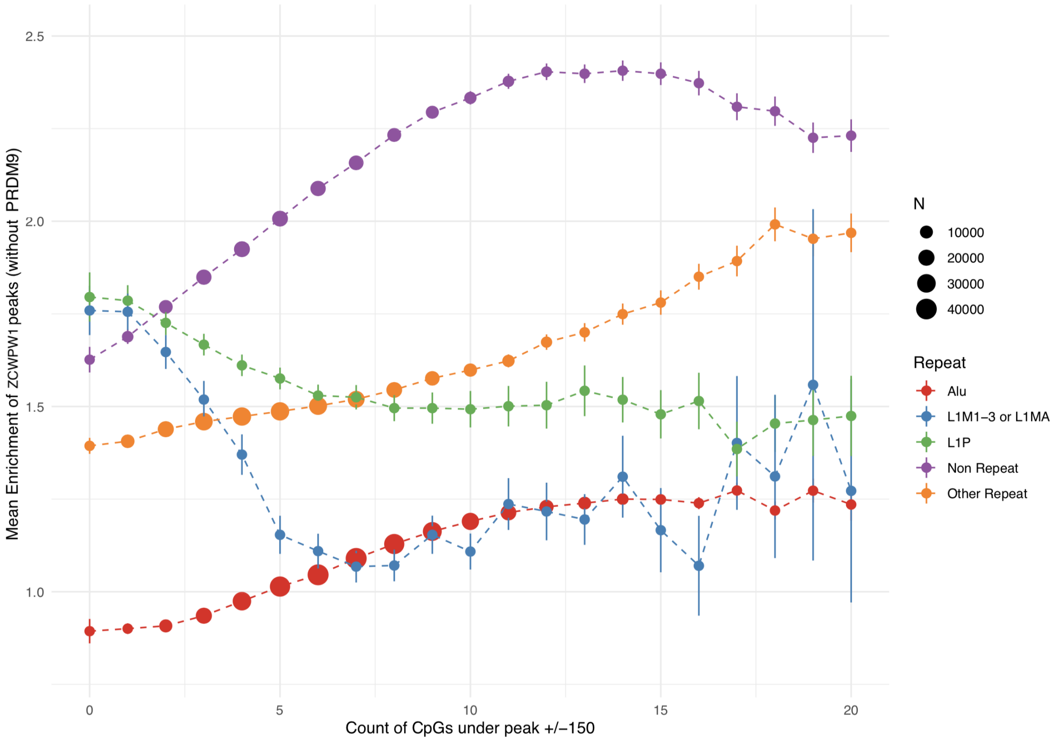


**Supplementary Figure 22.** CpG count around ZCWPW1 peaks (+/-150bp, with input coverage>5) is positively associated with ZCWPW1 enrichment in both peaks overlapping Alus and peaks not overlapping Alus, but not at L1M1-3, L1MA or L1P repeats. Error bars show ±2 s.e. of the mean.

**Supplementary Table 2. Breeding performance of *Zcwpw1*$^{-/-}$ females**

| Mouse | Genotype | Litter | Age at time of litter/death (months) | Pups born | Cumulative pups born | Frequency (time since previous litter in weeks/days) | Average number of pups/litter over breeding life |
|---|---|---|---|---|---|---|---|
| 1 | *Zcwpw1*$^{-/-}$ | 1 | 3 | 9 | 9 | N/A | 4.1 |
| | | 2 | 4 | 7 | 16 | 3w 5d | |
| | | 3 | 5 | 4 | 20 | 3w 2d | |
| | | 4 | 5.75 | 4 | 24 | 3w 1d | |
| | | 5 | 6.5 | TLL* | 24 | 3w 1d | |
| | | 6 | 7.5 | 3 | 27 | 4w 0d | |
| | | 7 | 8.25 | 2 | 29 | 3w 2d | |
| 2 | *Zcwpw1*$^{-/-}$ | 1 | 2.75 | 7 | 7 | N/A | 7 |
| | | 2 | 3.5 | 5 | 12 | 3w 3d | |
| | | 3 | 6 | 12 | 24 | 9w 6d | |
| | | 4 | 7 | 4 | 28 | 4w 1d | |
| | female culled | | 9 | N/A | 28 | 8w 3d | |

All females were crossed with a WT male. N/A, not applicable; TLL, total litter loss; *assigned a value of 0 as the number of pups born in the total counts.

**Supplementary Table 3. Fertility measures in WT, *Zcwpw1*$^{-/-}$ and *Prdm9*$^{-/-}$ males**

| Mouse | Genotype | Age at death (weeks, days) | Total body weight (g) | Lean body weight (g) | Paired testes weight (mg) | Paired testes weight / g of total weight | Paired testes weight / g of lean weight | Sperm count in millions / paired epididymides (average of 4 counts) |
|---|---|---|---|---|---|---|---|---|
| 1 | +/+ | 10w 1d | 26.2 | 19.8 | 179.2 | 6.84 | 9.05 | 20.45 |
| 2 | +/+ | 10w 1d | 27.4 | 20.42 | 183.8 | 6.71 | 9 | 32.5 |
| 3 | +/+ | 9w 3d | 28.6 | 22.56 | 190.4 | 6.66 | 8.44 | 25.95 |
| 4 | *Zcwpw1*$^{-/-}$ | 9w 5d | 26.7 | 20.66 | 49.8 | 1.87 | 2.41 | 0 |
| 5 | *Zcwpw1*$^{-/-}$ | 9w 6d | 25.8 | 19.7 | 35 | 1.36 | 1.78 | 0 |
| 6 | *Zcwpw1*$^{-/-}$ | 8w 5d | 27.9 | 21.98 | 56.9 | 2.04 | 2.59 | 0 |
| 7 | *Zcwpw1*$^{-/-}$ | 9w 5d | 25.1 | 18.89 | 39.4 | 1.57 | 2.09 | 0 |
| 8 | *Zcwpw1*$^{-/-}$ | 9w 5d | 23.2 | 19.62 | 50.9 | 2.19 | 2.59 | 0 |
| 9 | *Prdm9*$^{-/-}$ | 12w 4d | 28.3 | 21.77 | 50.3 | 1.78 | 2.31 | 0 |
| 10 | *Prdm9*$^{-/-}$ | 12w 5d | 27.9 | 22.36 | 49.2 | 1.76 | 2.2 | 0 |

Fertility was assessed in mice ranging from 8 to 12 weeks of age through measurement of paired testes weight and sperm count.

**Supplementary Table 4. Impaired synapsis in *Zcwpw1*$^{-/-}$ males**

| Mouse | Genotype | Type of synaptic error | | | Normal cells | Cells analysed | % synapsis |
|---|---|---|---|---|---|---|---|
| | | Tangled | Multibody | Split XY | | | |
| 1 | +/+ | 0 | 1 | 0 | 50 | 51 | 98 |
| 2 | +/+ | 1 | 1 | 1 | 49 | 52 | 94 |
| 3 | +/+ | 0 | 3 | 0 | 53 | 56 | 94.6 |
| 4 | *Zcwpw1*$^{-/-}$ | 1 | 49 | 0 | 1 | 51 | 2 |
| 5 | *Zcwpw1*$^{-/-}$ | 17 | 34 | 0 | 0 | 51 | 0 |
| 6 | *Zcwpw1*$^{-/-}$ | 20 | 31 | 1 | 1 | 53 | 1.9 |
| 7 | *Prdm9*$^{-/-}$ | 31 | 21 | 0 | 3 | 55 | 5.4 |
| 8 | *Prdm9*$^{-/-}$ | 31 | 15 | 0 | 3 | 49 | 6 |

The number of normal pachytene cells showing full synapsis of all autosomes and sex chromosomes (expressed as "% synapsis" of all cells analysed) was determined by immunostaining of testis chromosome spreads against SYCP3, HORMAD2 and γ-H2AX (see images in **Supplementary Figure 5**). The nature of the defects observed in cells with asynapsis was recorded as either "tangled" (when chromosomes pair with the wrong partner, forming branched tangled structures); "multibodies" strongly positive for HORMAD2 (when asynapsed chromosomes form multiple XY-like bodies which end up merging with each other, and with the XY body; or "split XY" (when the X and Y sex chromosomes are found away from each other in different areas of the cell nucleus).
